# Defect-Facilitated Buckling in Supercoiled Double-Helix DNA

**DOI:** 10.1101/259689

**Authors:** Sumitabha Brahmachari, Andrew Dittmore, Yasuharu Takagi, Keir C. Neuman, John F. Marko

**Affiliations:** Department of Physics and Astronomy, Northwestern University, Evanston, IL 60208, USA; Laboratory of Single Molecule Biophysics, National Heart, Lung,and Blood Institute, National Institutes of Health, Bethesda, MD 20892, USA; Department of Molecular Biosciences, Northwestern University, Evanston, IL 60208, USA

## Abstract

We present a statistical-mechanical model for stretched twisted double-helix DNA, where thermal fluctuations are treated explicitly from a Hamiltonian without using any scaling hypotheses. Our model applied to defect-free supercoiled DNA describes coexistence of multiple plectoneme domains in long DNA molecules at physiological salt concentrations (≈ 0.1 M Na^+^) and stretching forces (≈ 1 pN). We find higher (lower) number of domains at lower (higher) ionic strengths and stretching forces, in accord with experimental observations. We use our model to study the effect of an immobile point defect on the DNA contour that allows a localized kink. The degree of the kink is controlled by the defect size, such that a larger defect further reduces the bending energy of the defect-facilitated kinked end loop. We find that a defect can spatially pin a plectoneme domain via nucleation of a kinked end loop, in accord with experiments and simulations. Our model explains previously-reported magnetic tweezer experiments [1] showing two buckling signatures: buckling and ‘rebuckling’ in supercoiled DNA with a base-unpaired region. Comparing with experiments, we find that under 1 pN force, a kinked end loop nucleated at a base-mismatched site reduces the bending energy by ≈ 0.7 *k*_B_*T* per unpaired base. Our model predicts coexistence of three states at the buckling and rebuckling transitions that warrants new experiments.

## I. INTRODUCTION

Double-helix DNA is a semiflexible polymer (wormlike chain) with a bending persistence length *A* ≈ 50 nm [2–4]. The stacked double-helical structure of DNA allows only a few degrees of bend at the base-pair (bp) length scale (double-helix DNA has 0.34 nm/bp); however, thermally-generated bends correlated over hundreds of base pairs lead to large bending deformations. This results in a persistence length which is much larger than the length scale associated with the chemical building blocks, i.e., base pairs. In this article, we study the influence of a defect at the base-pair length scale, such as a base-mismatched region [1, 5] or a permanent DNA kink [6] on the statistical-mechanical properties of many persistence lengths-long double helices.

The double-helical structure also imparts a torsional rigidity to DNA, with a twist persistence length *C* ≈ 95 nm [7–9]. The two helically-linked strands of the double helix have a *linking number* ∆Lk ≡ Lk – L/h, where *L* is the total DNA contour length, *h* ≈ 3.6 nm is the intrinsic helix-repeat length, and Lk is the net linking of the two base-paired strands. The zero linking number state (∆Lk = 0) corresponds to a relaxed double helix. Twisting the double helix perturbs the linking number resulting in a buildup of DNA torque.

Under a constrained topology or fixed linking number condition, twisted DNA can buckle into a superhelical structure called a *plectoneme* [Fig. 1(a)]. The *writhe* linking number contribution, a geometric invariant, associated with the plectoneme structure determines the equilibrium state of twisted or *supercoiled* DNA. DNA supercoiling is associated with various cellular functions, such as genome organization, gene expression, and DNA recombination [10–13]. Bacterial DNA is maintained in a supercoiled condition, consequently plectonemes are a common occurrence in the cell. In *vivo* manipulation of DNA linking number, carried out by topoisomerase enzymes [14], is essential for topological simplification of the entangled state of the genome.

**FIG. 1. (a).**
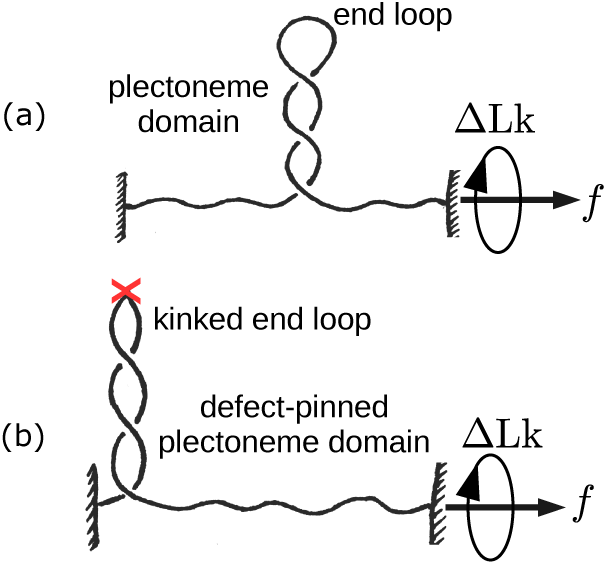
Schematic of stretched defect-free double-helix DNA plectonemically buckled under torsional stress. A plectoneme domain contains an end loop that is associated with a nucleation cost of the domain. (b) DNA with a defect located on its contour (denoted by an ‘X’). The defect allows a localized DNA kink that favors nucleation of a *defect-pinned* plectoneme domain characterized by an energy-saving kinked end loop. However, the immobile nature of the defect spatially pins the domain costing diffusion entropy.

Single molecule experiments have been crucial in studying DNA mechanical response to linking number perturbation. While some experiments have directly visualized plectonemes using electron microscopy [15] and DNAs with fluorescent labels [16]; others have studied plectonemic buckling using tweezer techniques [6–9, 16–18]. Magnetic or optical tweezers allow precise control of DNA linking number under an applied stretching force [Fig. 1]. Theoretical models have also played an indispensable role in understanding the mechanical behavior of DNAs and the plectonemic state [19–25]. While the previous theoretical work have greatly enhanced our understanding of DNA supercoiling, there are still poorly understood aspects, such as whether DNA buckles to form a single or multiple plectonemic domains.

The double helix is not a homogeneous polymer, the elastic moduli associated with an AT base pair may be different from that of a GC. Should the DNA thermal persistence length be sequence dependent? For a double helix containing random base pairs, a simplified view of a constant bending and twisting moduli suffices. However, for DNA fragments containing certain periodic base pair arrays (e.g., a positioning sequence, such as the 601 sequence [26]), an intrinsic curvature in the double helix backbone reduces the local bending stiffness. Positioning sequences are thought to regulate cellular function via biasing nucleosome positioning and altering genome access for transcription factors and other DNA-binding proteins [27–29]. DNA containing a positioning sequence *spatially-pinned defect*. The thermal persistence length associated with the defect is smaller, which reduces the energy cost of a localized bend. Such a defect on the double-helix contour can arise from a variety of other sources as well, such as a base-mismatched region [1, 5], a DNA hairpin [6], a single-stranded DNA bulge, or a protein-induced DNA kink [30, 31]. The defect, however, suppresses diffusion of the bent structure which decreases entropy. Experiments [1, 5] and simulations [32] studying supercoiling DNA with unpaired bases have shown that a defect can spatially pin a plectoneme domain.

In this article, we describe a twistable worm-like chain model for supercoiled DNA that agrees with experimental observations. We model thermal fluctuations in plectonemic DNA as a perturbation around its mean-field structure that improves upon the scaling-like treatment employed in previous work [19–25]. We use our model to study supercoiling of DNA with an immobile defect, where the energy cost is reduced for a bend localized at the defect site [Fig. 1(b)].

Motivated by recent experiments on supercoiling DNA with base-pair mismatches [1], we consider the possibility of an immobile DNA *kink* at the defect location that may spatially pin a plectoneme domain. Experiments showed a novel second buckling signature or ‘rebuckling’ transition that is derived from the location of the defect on the DNA contour [1]. Our model reproduces the rebuckling transition, explains the thermodynamic picture underlying existing experiments [1], and makes predictions for future experiments.

*Outline of this paper*. Sec. II contains theory [Sec. II A] and numerical results [Sec. II B] for supercoiled defect-free DNA, where we also compare with available experimental data. We analyze the effects of salt concentration and length of the supercoiled DNA molecules on its statistical mechanics [Secs. IIB 1 and IIB2]. Our results also explain the abrupt buckling transition and coexistence of multiple buckled domains in supercoiled DNA.

Sec. III describes how the model can take into account an immobile point defect on the DNA [Sec. III A], and explains the results of its numerical solution [Sec. III B]. Our results reproduce the second buckling signature or rebuckling transition observed previously in magnetic tweezers experiments [1] [Sec. III B], and explains the free energy picture underlying buckling [Sec. III B 1] and rebuckling transitions [Sec. III B2]. We highlight the role of the size of the defect and predict coexistence of multiple states at the transitions [Sec. III B 3]. In Sec. IIIC, we compare our theoretical results with experimental data of Ref. [1]. We explain the observed shift in transition points [Sec. III C 1] and quantitatively connect our theoretical defect size parameter with experiments [Sec. IIIC 2]. The experimental trends for extension jump with varying defect sizes [Sec. III C 3], and the force and salt dependence of the rebuckling signal [Sec. III C4] are also in accord with the theoretical results. Finally, in Sec. IV, we conclude with a summary and discussion of future prospects.

## II. SUPERCOILED DEFECT-FREE DNA

We consider DNA as a charged semiflexible polyelectrolyte with bending persistence length *A =* 50 nm and twist persistence length *C* = 95 nm. We define *β*^-1^ ≡ *k*_B_*T* and use *T =* 290 K for all numerical purposes. In the following, we investigate the mechanical response of a double-helix DNA stretched under a constant force f and subject to twist such that the change in the DNA linking number is ∆Lk. We present the model schematically in this section; mathematical details are in Appendices A and B.

### A. Model

We partition total DNA length (*L*) and linking number (∆Lk) into a force-extended or unbuckled state *(L_u_*, Lk*_u_*), a plectonemically-buckled superhelical state *(L_p_*, Lk*_p_*), and *m* end loops each of size γ corresponding to *m* plectoneme domains:

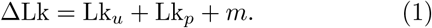

We consider the writhe linking number contribution from each end loop to be ≈ 1. There is also a constraint of fixed total DNA length: *L = L_u_* + *L_p_* + mγ.

The total free energy of a stretched-twisted DNA is written as follows.

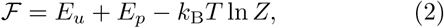

where *E_u_* corresponds to the force-extended or unbuckled part of the DNA containing contributions from DNA twist and force extension [Eq. (B.3)]. *E_p_* is the mean-field energy corresponding to the plectonemically-buckled state [Eq. (B.1)]; and *—k*_B_*T ln Z is the* free energy contribution from thermal fluctuations of the DNA [Eq. (A.7)]. The total free energy is minimized with respect to partition of linking number to obtain the equilibrium linking numbers corresponding to the force-extended and plectoneme states.

#### 1. Buckled state

##### a. Plectoneme superhelix

The plectoneme state is characterized by superhelically-coiled DNA, and has no force-extension energy [Fig. 1(a)]. However, transverse fluctuations of the DNA within the superhelical structure are controlled by both the applied tension and electrostatic interactions (Appendix A). The plectoneme state costs bending energy, but at the same time reduces twist energy due to the writhe linking number contribution associated with the superhelical structure. Following White’s theorem of partition of linking number into twist and writhe [33], the linking number in the plectoneme state Lk*_p_* is divided as follows:

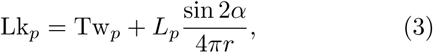

where the first RHS term is the twist linking number contribution, and the second term is the total writhe of a helical structure with opening angle α, helical radius r, and total plectoneme length *L_p_* [19, 33].

##### b. Plectoneme end loop

The end loop is a finite-sized DNA structure where the double helix bends back in a plectoneme [Fig. 1(a)]. We compute the equilibrium size of the end loop: 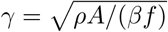, via separately minimizing the associated elastic energy expense [23, 34]:

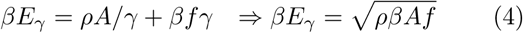

The parameter *ρ* depends on the geometry of the loop, such that it is *2*π*^2^* for a circular loop, ≈ 14.0 for a “teardrop”-shaped loop with free ends [35], and ≈ 15.3 (exact) or ≈ 15.7 (simpler calculation) for an end-constrained teardrop loop [36, 37]. We use *ρ =* 16, however, we note that a small change (≈ 10%) in the numerical value of *ρ* does not alter our conclusions. In the later part of the paper concerning defects, we analyze the effect of a relative variation in *ρ* [Eq. (8)].

#### 2. Thermal fluctuations

The mean-field structure of the force-extended state is a twisted straight line, whereas, that of the plectoneme state is a regular helix made up of self-writhed twisted DNA. At a finite temperature, we treat thermal fluctuations as a small perturbation around the mean-field structure (Appendix A).

The total energy associated with transverse fluctuations of the DNA about its mean-field shape is as follows [Eq. (B.4)].

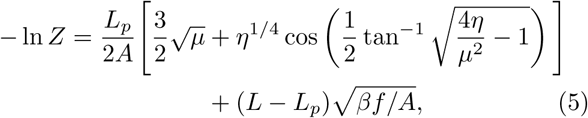

where *µ ≡* β *Af cos α*, is the effective tension in each of the two helically wrapped strands of the plectoneme; and 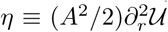, is the effective electrostatic modulus of uniform radial deformations in the plectonemic superhelix (see Appendix A).

The first bracketed term in Eq. (5) corresponds to thermal fluctuations in the plectoneme structure [Eq. (A.7)]. Note that there are four degrees of freedom associated with transverse fluctuations in a plectoneme structure, two for each of the plectonemic strands. In a conveniently chosen reference frame [Eq. (A.3)], three of the degrees of freedom fluctuate independently under the external tension, as seen in the first term inside the brackets (the term proportional to 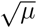). The second term within the brackets, dependent on the strength of the electrostatic repulsions via *η*, corresponds to electrostatically-coupled fluctuations where the two strands displace relative to each other. The second term in Eq. (5) corresponds to tension-controlled transverse fluctuations in the force-extended part of the DNA.

We note that this part of the computation substantially improves on the prior work where a scaling-like free energy cost from cylindrical confinement is used to account for fluctuations in the plectonemic superhelix [4, 21, 25, 38, 39]. Our approach, resulting in Eq. (5), proposes an explicit computation of thermal fluctuations, treating them as a perturbation about the mean-field helical geometry of the plectoneme. Eq. (5) is consistent with the previously assumed scaling: 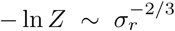 (where *σ_r_* is the fluctuation in plectoneme radii [Eq. (A.8)]), but does not depend on any unknown scaling constants.

Note that a similar treatment of thermal fluctuations in two helically-intertwined DNAs or braids can be found in Ref. [34]. DNA braids have the same geometry as plectonemes, however experimentally studied braids [40–42] are force extended, while plectonemes are buckled structures that do not have force-extension energy [Appendix A].

#### 3. Extension and torque

DNA extension *z* can be obtained from the negative force derivative of the total free energy [Eq. (2)]. Hence, DNA lengths in the force-extended and buckled states are respectively associated with positive and zero contributions to extension. Thermal fluctuations further reduce DNA extension, which is a sub-leading order effect.

Torque in the DNA *τ* is obtained via linking number derivative of the total free energy. The free energy is harmonic in twist linking number, however, conversion of twist to writhe in the plectoneme-coexistence state influences the torque response.

##### a. Partition function

We sum over states containing all possible lengths *(L_p_)*, and numbers of domains (*m*) of plectoneme to construct a canonical partition function:

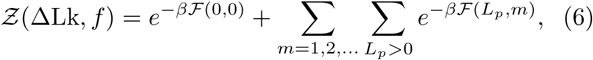

where the coexistence-state energies **F**(*L_p_*, m) [Eq. (2)] are obtained by free-energy minimization, which also ensures balance of torque in the summed-over states. Energy minimization for a coexistence state with fixed *L_p_* and m determines the equilibrium plectoneme radius (*r*) and opening angle *(α).*

Equilibrium values of end-to-end extension (*z*), number of plectoneme domains (*m*), torque in the DNA (*τ*), and the total plectoneme length *(L_p_)* at a fixed force and fixed linking number are obtained from the partition function [Eq. (B.5)].

##### b. Probability distributions

In a canonical ensemble of fixed force f and fixed linking number ∆Lk, both *z* and τ undergo equilibrium fluctuations. The total probability distribution of X ∈ {*z,τ*} is obtained by adding the contributions from various states in the partition sum:

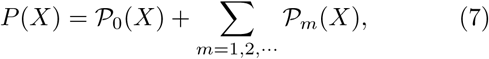

where 𝒫_0_ corresponds to the force-extended state, and 𝒫*_m_* (where m ∈ {1, 2, · · ·}) is the contribution from the buckled state featuring coexistence of m plectoneme domain(s). States corresponding to different plectoneme lengths in the partition sum are already taken into account in 𝒫*_m_* [Eq. (B.6)].

#### B. Results: Defect-Free DNA

In this section, we describe the numerical solutions of the model and compare our results with experimental data.

##### 1. Supercoiling at physiological salt

We begin with a discussion of rather short DNA molecules (2 kb), subject to twist under physiological salt conditions (≈ 0.15 M Na^+^).

###### a. Slightly twisted DNA: unbuckled state

Untwisted DNA extension is 80–90% of its total contour length under 0.5–2 pN stretching force [Fig. 2(a)]. Higher stretching forces suppress the excursions of the DNA lateral to the force direction, resulting in a longer extension at zero linking number. Small twisting of the double helix results in a linear buildup of DNA torque [Fig. 2(b)]. Change in extension upon slight twisting of the double helix is small. This is due to twist-induced chiral fluctuations in DNA that leads to partial twist screening, which is taken into account via a renormalized twist stiffness for stretched-unbuckled DNA [Eq. (B.3)] [20].

**FIG. 2.**
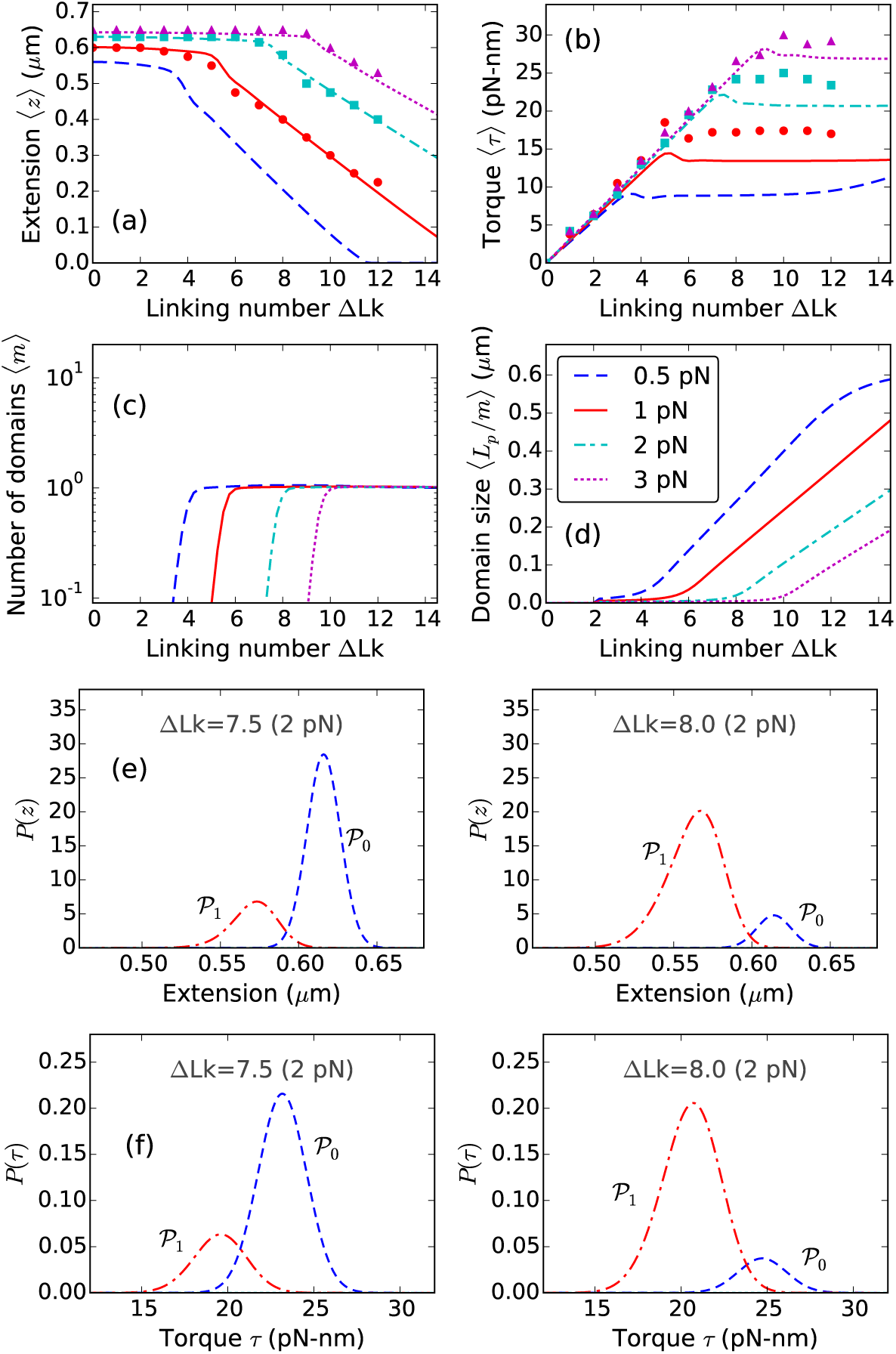
Theoretical curves for supercoiled 0.7 μm (≈2 kb) DNA molecule, stretched under 0.5 (blue dashed lines), 1 (red solid lines), 2 (cyan dot-dashed lines), and 3 pN (magenta dotted lines) applied forces at 0.15 M Na^+^. Experimental data reproduced from Ref. [8] are plotted for 1 (red circles), 2 (cyan squares), and 3 pN (magenta triangles), (a) Extension versus linking number, shows a flat unbuckled regime at lower linking numbers. Extension decreases steeply at higher linking numbers corresponding to coexistence of a plectoneme state. The extension discontinuity connecting the two slopes corresponds to the buckling transition, (b) Torque increases linearly in the unbuckled state, and then saturates as a part of the DNA buckles to form plectoneme. Constant torque in the plectoneme coexistence state is due to the writhe contribution of the plectoneme geometry that screens DNA twist. The small overshoot in torque as well as the discontinuity in extension near the buckling transition point is related to the end loop-introduced nucleation cost of a plectoneme domain, (c) Equilibrium number of plectoneme domains grows to unity at the buckling transition point, showing nucleation of a plectoneme domain, (d) Average plectoneme domain size increases after the buckling point, indicating addition of superhelical turns to the buckled domain. Probability density of (e) extension and (f) torque near the buckling point is bimodal [ΔLk=7.5 and 8.0 under 2 pN force]. The modes of the distributions at higher extension and torque correspond to the unbuckled state (*<*scP*>o*, blue dashed lines); whereas, the lower extension and torque modes correspond to the one-domain plectoneme state (*<*scP*>*_1_, red dot-dashed lines). The probability distributions show an increase in the average occupancy of the buckled state (*<*scP*>*_1_) as the linking number is increased near the buckling point.

Linear torque indicates a constant twist stiffness in DNA, which is a consequence of the rigidly-stacked double-helical structure. A softer structure, two-intertwined nicked DNAs or a “braid” has a linking number dependent twist stiffness [34].

###### b. Buckled DNA

At higher linking numbers, the double-helix buckles into a self-writhed plectoneme struc ture [4, 15]. Plectonemes with higher pitch-to-radius ratio have a substantial *writhe* linking number density [Eq. (3)], as a result, buckling avails conversion of twist into writhe. Post-buckling torque is nearly constant [Fig. 2(b)] suggesting that increasing the linking number in buckled DNA does not increase DNA twist but increases total writhe. Plectonemes save DNA twist energy but cost bending energy, hence buckling is favored only above a critical torque that corresponds to a *critical linking number.* Higher stretching forces stabilize the unbuckled state, resulting in an increase in the critical linking number [Fig. 2]. The mean-field plectoneme state does not contribute to end-to-end extension, resulting in a steeper decrease in extension in the coexistence regime [Fig. 2(a)].

###### c. Plectoneme domains

Figure 2(c) shows the appearance of a plectoneme domain at the buckling transition, which grows in size as the linking number is increased beyond the critical value [Fig. 2(d)]. The increase in the average size of the plectoneme domain is due to equilibrium DNA length being passed from the unbuckled into the buckled state, which increases the number of superhelical turns in the plectoneme domain.

###### d. Abrupt buckling

The buckling transition is abrupt due to the finite-sized end loop that associates a *nucleation cost* to a plectoneme domain. The discontinuity in extension and the overshoot in torque at the buckling point [Fig. 2(a)-(b)] characterizes the abrupt nature of the transition. Near the buckling point, the unbuckled and the plectoneme states are thermally accessible, which implies that the probability of occupancy of either state is non-zero. Figure 2(e) shows bimodal probability densities of extension near the buckling point, where the unbuckled and the plectoneme states correspond to the higher (𝒫0) and lower (𝒫1) extension modes, respectively. The discontinuity in extension at the buckling point (i.e., the non-zero distance between the two extension modes) is due to the fact that a buckled domain cannot be smaller than an end loop, which is *O*(1) DNA persistence length in size. As the linking number is increased at the buckling point, the probability of occupancy of the buckled state increases, and that of the unbuckled state decreases [Fig. 2(e)].

The average torques in the two fluctuation-accessible states (*<*scP*>*_0_ and *<*scP*>*_1_) at the buckling transition are different; this is due to the writhe associated with the nucleated buckled domain (plectoneme and the end loop). Figure 2(f) shows the bimodal torque distributions near the buckling point. Equilibrium fluctuations between the two states of different torques: *<*scP*>*_0_ and *<*scP*>*_1_, lead to an overshoot behavior seen in the ensemble-averaged DNA torque at the buckling transition [Fig. 2(b)]. A nonmonotonic mechanical torque, i.e., decreasing torque with increasing linking number indicates negative torsional stiffness, a signature of mechanical instability. However, the ensemble-averaged torque may show nonmonotonic behavior in an equilibrated system with monotonic mechanical torque. This is a consequence of nonmonotonic behavior of *torque fluctuations* near the buckling transition (see Sec. V of Ref. [23]). There is experimental evidence of an overshoot in DNA torque at the buckling transition [8, 18].

###### e. Coexistence of multiple plectoneme domains in longer DNA molecules

Entropic stabilization of plec-toneme domains from one-dimensional diffusion along the DNA contour and exchange of DNA length among domains (i.e., fluctuations in relative size of the domains) increases logarithmically with DNA length [Eq. (B.1)]

[23]. This leads to proliferation of multiple domains in supercoiled long DNA molecules (≲10 kb). Diffusion of plectonemes and coexistence of multiple domains have been observed in DNA visualization experiments [5, 16].

Figure 3 shows buckling behavior in long DNA molecules. The critical linking number is an extensive quantity that increases with DNA length [Figs. 2(a) and 3(a)]; however, the critical buckling torque is intensive and remains roughly the same for different length molecules [Figs. 2(b) and 3(b)].

**FIG. 3.**
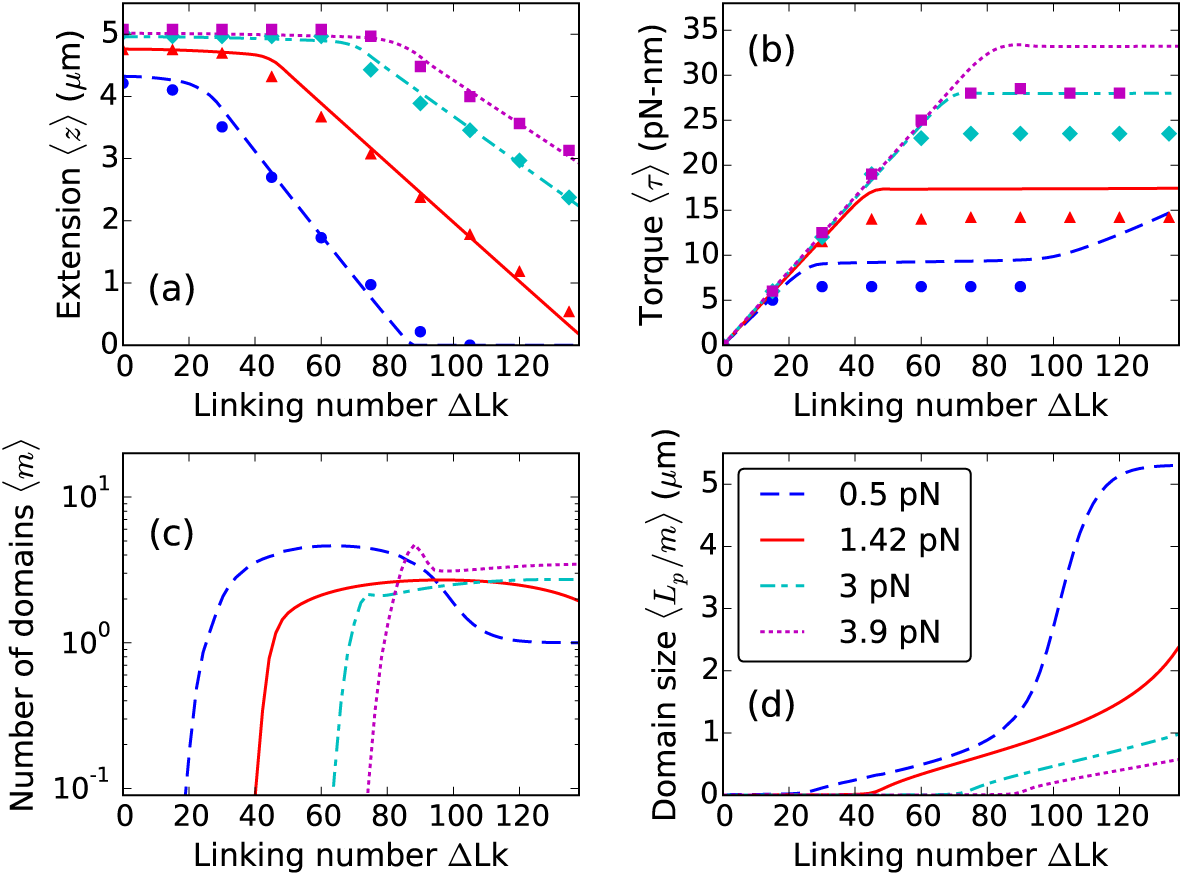
Theoretical curves for supercoiled 5.4 *μ*m (≈ 16 kb) DNA, stretched under 0.5 (blue dashed lines), 1.42 (red solid lines), 3 (cyan dot-dashed lines), and 3.9 pN (magenta dotted lines) forces at 0.1 M Na^+^. Experimental data for 0.5 (blue circles), 1.42 (red triangles), 3 (cyan diamonds), and 3.9 pN (magenta squares) are reproduced from Ref. [9]. (a) Extension and (b) Torque plotted as a function of linking number show twisting behavior at lower linking numbers and plectoneme buckling at higher linking numbers, (c) Equilibrium number of plectoneme domains show proliferation of multiple plectoneme domains in the coexistence state. At higher forces, long molecules show a non-monotonic increase in the number of plectoneme domains at the buckling transition due to the large entropy associated with plectoneme diffusion. However, in the purely-plectoneme state (i.e., the zero extension state, refer to the 0.5 pN case, at linking numbers ≳ 90), high stability of plectoneme superhelices and absence of diffusion entropy results in favoring a single plectoneme domain. Torque in the purely-plectoneme state increases because the DNA twist increases, (d) The steepness in the increase of the average domain size increases in the purely-plectoneme state due to coalescence of plectoneme domains.

The larger configuration entropy associated with long DNA molecules reduces the nucleation energy of a plectoneme domain, resulting in proliferation of multiple domains in the buckled state [Fig. 3(c)]. At lower forces, the nucleation energy cost is further reduced resulting in an increased tendency to proliferate new plectoneme domains. However, in the *purely-plectoneme state* (i.e., the zero-extension state, where the entire DNA is in the plectoneme state, L*_u_* = 0), a single plectoneme domain is favored due to reduced diffusion entropy of a domain. Figure 3(c) shows coalescence of multiple domains as the linking number is increased in the purely-plectoneme state, which is the result of a highly stable plectoneme superhelix compared to an end loop.

Energy of the unbuckled and the plectoneme states vary linearly with force, whereas, that of the end loops vary as the square root of the applied stretching force. This leads to an increased probability of small plectoneme domains (i.e., end loop with a minimal amount of plectoneme superhelix) at the buckling transition under higher forces. As a result, twisted long DNA, under higher forces, buckles via nucleation of a few small loops that coalesce at a slightly higher linking number due to increased stability of plectonemic superhelices. This is seen as a small overshoot in the number of plectoneme domains at the buckling transition under higher forces [Fig. 3(c)].

Post-buckling torque is mostly constant in the plectoneme-coexistence state, however, increasing the linking number in the purely-plectoneme state causes an increase in the DNA twist, reflected in an increase in the torque [Fig. 3(b)].

Theoretical post-buckling torque appears to be an underestimation compared to experiments of Ref. [8] [Fig. 2(b)], where torque was inferred from angular fluctuations of an optically-trapped DNA-tethering particle. On the other hand, torque reported in Ref. [9], derived from the slope of the extension curves using Maxwell relations [43] is lower than the theoretical values [Fig. 3(b)]. While in-situ torque measurement is a remarkable step forward for experiments, equilibration might be an issue. Maxwell relations are a robust tool for torque estimation, however, the procedure employed by Ref. [9] assumes a constant torque in the plectoneme-coexistence state. Our model, although devoid of the above issues, assumes a regular plectoneme geometry and ignores any energy contribution from distortion of the helical plectoneme structure. Numerical values of the DNA effective charge [21, 44] as a function of salt concentration also affects the coexistence state torque. These small discrepancies call for future attention both from the theoretical and experimental perspectives.

#### 2. Effect of salt concentration

An increased ionic concentration of the solution strengthens the electrostatic screening of the charges on the DNA backbone, which reduces the effective excluded diameter of DNA (measured in Debye-Hückel screening length *λ_D_*). At lower salt concentrations, a larger screening length mimics stronger self-avoidance in DNA, which shifts the free energy balance in favor of looped structures over plectonemic superhelices. This effectively translates into a higher tendency to proliferate multiple domains of plectoneme as the salt concentration is lowered.

##### a. Buckling transition

The critical linking number increases with a decrease in the salt concentration (Fig. 4). Larger excluded diameter of DNA at lower salt concentrations increases the bending energy of the plectoneme state, leading to an increase in the critical linking number. The post-buckling state is that of many domains at lower salts (≈ 0.01 M Na^+^) (Fig. 4), which is again related to the increased energy of plectoneme superhelices that favors nucleation of many small plectonemes instead of a single large domain.

**FIG. 4.**
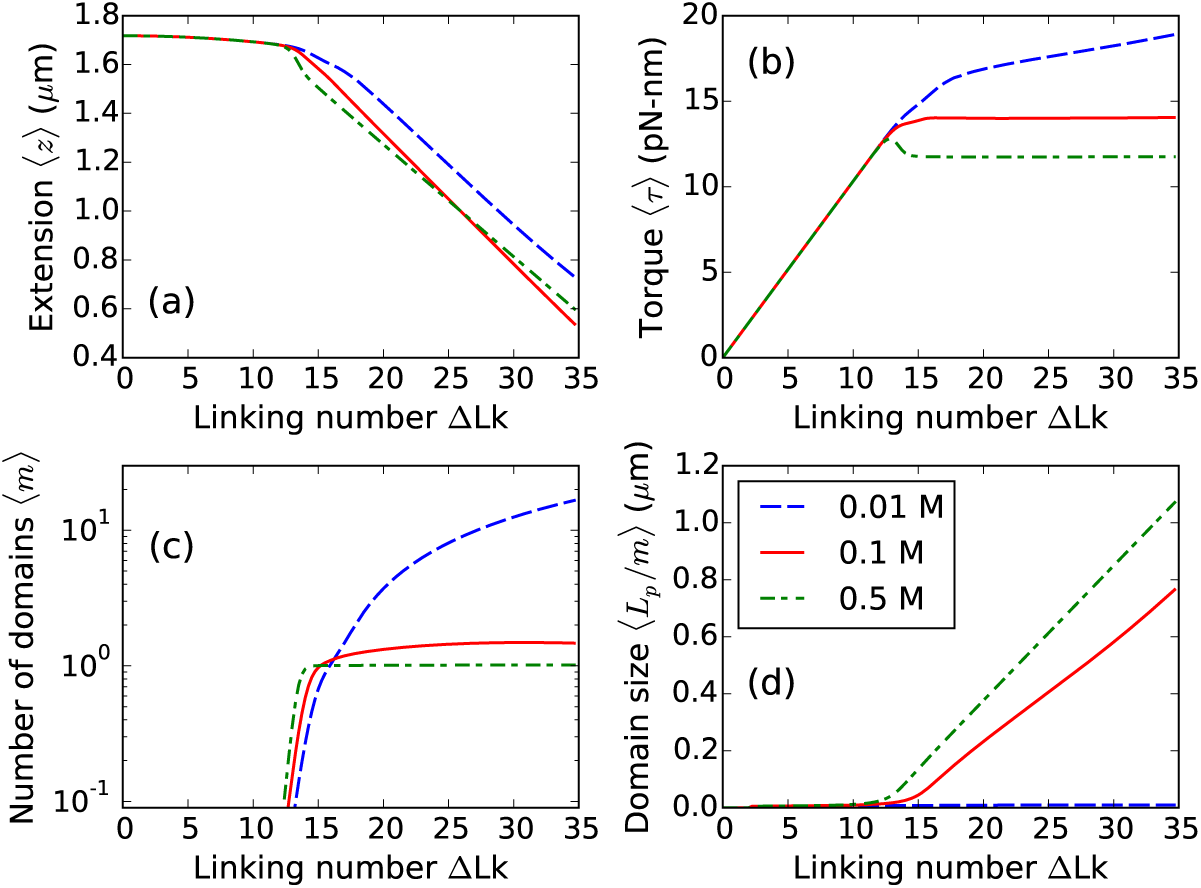
Effect of salt concentration on defect-free DNA. Supercoiled 2*μ*m DNA at 1 pN stretching force under 0.01 (blue dashed line), 0.1 (red solid line), and 0.5 M Na^+^ (green dot-dashed line), (a) Extension and (b) Torque shows a more rounded buckling transition for lower salts due to lower stability of plectoneme superhelices. Note that the torque increases in the buckled state for lower salts corresponding to increase of DNA twist due to less twist screening by unstable plectoneme superhelices. (c) Number of plectoneme domains increase in the buckled state for lower salts, whereas, the buckled state is constituted of a single plectoneme domain at higher salts, (d) Average size of a domain increases in the higher salt case. For lower salts, proliferation of multiple domains lead to a very small domain size in the buckled state.

##### b. Multidomain plectoneme for lower salts

Torque in the buckled state is flat for higher salts and increases slowly with linking number for lower salt concentrations (Fig. 4). For lower salts, decreased stability of the plectonemic superhelix (due to higher DNA excluded diameter) causes a small increase in DNA twist in the plec-toneme coexistence state. The extension distributions, at low salt concentrations (≈ 0.01 M Na^+^), are bimodal at the buckling transition and remain multimodal in the buckled state due to coexistence of multiple plectoneme domains (Fig. 5), which may be visible in tweezer experiments. However, increased fluctuations may decrease the resolution of the two peaks, producing a broad-tailed single Gaussian shape.

**FIG. 5.**
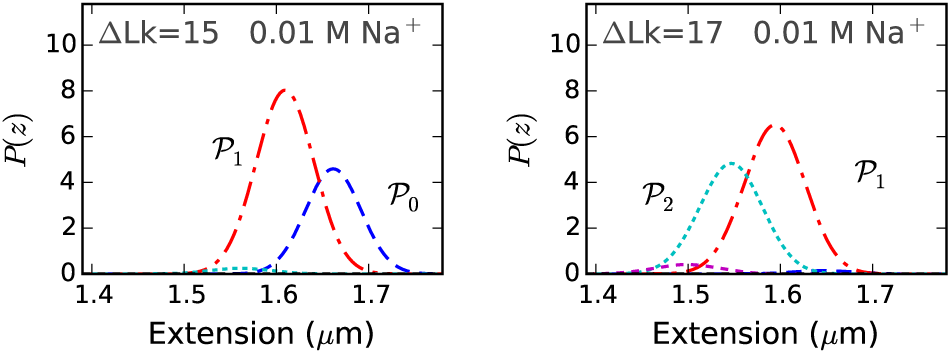
Probability density of extension at 0.01 M Na^+^ under 1 pN force near the buckling transition for a 2 µm DNA (≈ 6 kb) [Fig. 4]. The extension distribution is bimodal, however, the two modes, corresponding to unbuckled DNA (𝒫_0_, blue dashed line) and one plectoneme domain (𝒫_1_, red dot-dashed line), are less resolved at lower salts due to increased fluctuations. The probability distribution remains bimodal after the buckling point, due to appearance of multidomain plectoneme states (e.g., the two domain state 𝒫_2_, cyan dotted line at ∆Lk=17). Decreased stability of the plectoneme superhelix at lower salts result in coexistence of multiple plectoneme domains. Increasing the linking number in the buckled state increases the probability of occupancy of a plectoneme state with a larger number of domains.

DNA braids are structurally bulky, and as a result, mimic the low salt behavior of supercoiled DNAs. Braids show multimodal extension profiles corresponding to proliferation of buckled domains [34, 42].

We assume the unbuckled state to be decoupled from the electrostatics, because non-neighbor segments in unbuckled DNA are always distant. Nonetheless, increased repulsion between neighboring segments at lower salts (due to less DNA charge screening), is expected to induce additional stretching of the polymer. This maybe taken into account via a persistence length that gets longer with decreasing salt concentration. Experiments suggest a small change (≈ 10%) in the persistence length over a wide range of salt concentrations (0.01–1 M Na^+^) [45], which we ignore for simplicity. The ionic strength dependence of DNA torsional stiffness is also negligibly small [46].

##### c. Varied abruptness of the buckling transition

The discontinuous change in extension and overshoot in torque at the buckling point are measures of the abruptness of the transition, which decreases with decreasing salt concentration (Fig. 4). A more abrupt transition, i.e., two well-separated peaks in the extension profile, results from a larger size of the nucleated buckled domain. The nucleated domain at buckling consists of an end loop and a plectoneme comprising superhelical turns. When the superhelix is less stable, the case for lower salt concentrations (≈ 0.01 M Na^+^), the nucleated domain is an end loop with minimal superhelix, which leads to a less abrupt extension change at buckling.

When the salt concentration is increased, plectoneme-superhelical turns are increasingly stabilized, and the amount of superhelically wrapped DNA in the nucleated domain also increases. This produces a larger extension change (i.e., strongly bimodal extension distribution) at buckling for higher ionic strengths.

## III. DNA WITH A SPATIALLY-PINNED POINT DEFECT

In the following section, we describe an immobilized point defect on the DNA that can spatially-pin a kinked end loop. Consequently, nucleation of a spatially-pinned plectoneme domain may be favored at the defect site due to the higher bending energy of a teardrop end loop compared to a kinked loop. We introduce a defect size parameter *ε* that controls the relative stability of a kinked loop. We also predict a defect-size-dependent coexistence of three states at buckling and rebuckling transitions.

## A. Model for an Immobile Defect

We suppose that a defect acts as a soft-spot for bending deformations, such that the DNA can form a kink at the defect site [Fig. 1(b)]. We are motivated to describe defects arising from base-unpaired regions on the DNA [1], but the biological relevance of a defect-induced DNA kink is diverse, such as a protein-mediated DNA bend, or a single-stranded bulge on the DNA, or a damaged DNA base.

### Defect-pinned plectoneme domain

A plectoneme with its tip at the defect site can have a kinked end loop, thereby saving bending energy. However, the immobile nature of the defect restricts diffusion of a defect-pinned domain [Fig. 1(b)]. As a result, nucleation of a defect-pinned domain is expected to feature a competition between stabilization from lower bending energy of a kinked end loop and destabilization from spatial pinning.

### 1. Size of the defect ε

The defect may have a varied size that affects the degree of the defect-induced DNA kink; larger defects allow a sharper DNA kink, thus lowering the associated bending energy of a kinked end loop [37]. Following the scheme used for the defect-free DNA in Eq. (4), we use a defect-size dependent loop parameter: (1 – *ε*)^2^*ρ*, such that the energy of a kinked end loop 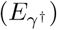 is a defect-dependent fraction of that of the teardrop loop:

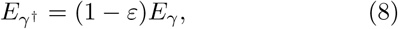

where 0 < *ε* < 1 is the size of the defect, and 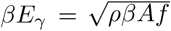 is the energy of a teardrop-shaped loop [Eq. (4)]. Energy minimization gives the equilibrium size of a defect-kinked end loop:

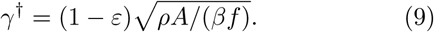

Higher values of *ε*, corresponding to a larger defect, stabilizes the defect-pinned loop by allowing a sharper kink at the defect site. The experimental counterpart of ε is the number of adjacent base-pair mismatches on a supercoiled DNA [1]. A defect with a larger number of unpaired bases further reduces the bending energy of a defect-kinked loop, which corresponds to a larger value of *ε.*

The total free energy of a stretched twisted DNA with a defect of size *ε* is given as:

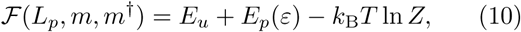

where the free energy of the coexistence state now depends on the length of the plectoneme *(L_p_)*, the number of mobile plectoneme domains (*m*), and the number of defect-pinned plectoneme domains (*m*^†^) which can be either 0 or 1 (i.e., *m*^†^ ∈ {0,1}). The size of the defect, ε affects the plectoneme state energy by changing the bending contribution associated with a kinked loop [Eq. (B.7)]. The total free energy is minimized for a fixed total length:

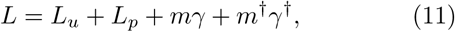

where γ and γ^†^ are the respective sizes of a teardrop and a kinked end loop. [Eqs. (4) and (9)]. The total linking number is also constrained, where both kinked and teardrop end loops contribute unit writhe linking number.

### 2. Critical size of the defect-pinned plectoneme

The size of a plectoneme domain is maximum at the end of the coexistence region, which is the purely-plectoneme state with no unbuckled DNA. However, a defect-pinned plectoneme can be made maximally big (corresponding to a *critical size)* at any point in the coexistence region by forcing its proximity to the tethering surface. For a defect located a distance *L\** from one of the ends, the defect-pinned plectoneme has a critical size of *2 L\** [1]. A defect-pinned domain must have the tip of the plectoneme at the defect site; this results in one of the ends of a critically big defect-pinned plectoneme domain coinciding with a surface-tether point of the DNA [Fig. 1(b)]. Consequently, at a coexistence point with total plectoneme length larger than *2 L\** there must be at least one mobile plectoneme domain (i.e., *m* ≥ 1).

#### a. Partition function

We sum over all possible sizes *(L_p_)* and numbers of the plectoneme domains, both mobile (m) and pinned (m^†^), to construct a canonical partition function:

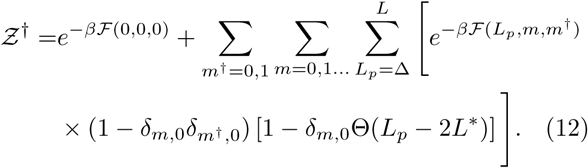

The above partition function imposes a critical size on the defect-pinned plectoneme via the product of the Kronecker delta with the Theta function [Eq. (B.8)]. The product of the two Kronecker delta functions ensures the presence of at least one end loop in the buckled state of the DNA. We get equilibrium values of observables such as extension, torque, and number of plectoneme domains from the above partition function [Eqs. (B.5)].

#### b. Extension distribution

The extension profile at a given linking number and force is also obtained from the partition function [Eq. (B.9)].

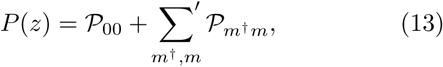

where the primed sum corresponds to a restricted sum as defined in the above partition function [Eq. (12)]. Here, 𝒫_00_ is the contribution from the unbuckled state; and *P_m_*_†_*_m_* is the contribution from a buckled state with m pinned and *m*^†^ mobile plectoneme domains. For instance, 𝒫_01_, 𝒫_10_, and 𝒫_11_ are the respective contributions from-the buckled state with one mobile plectoneme domain (*m*^†^ = 0 and *m* = 1), the buckled state with a defect-pinned domain *(m*^†^ *=* 1 and m = 0), and the two-domain plectoneme state containing one mobile and one defect-pinned domain (*m*^†^ = 1 and m = 1).

### B. Results: DNA with a Defect

Figure 6(a)-(b) shows extensions and torques, respectively, of a 6 kb DNA molecule with a defect of sizes *ε* = 0.05,0.15, and 0.3 as a function of linking number. Small twisting of the molecule leads to a small extension change and a linear torque buildup, as seen for defect-free DNA (Fig. 2). An increase in the DNA linking number leads to buckling of the double helix. The second buckling signature in the extension and torque curves [Fig. 6] or the *rebuckling transition* [1] is related to the critical size associated with the defect-pinned plectoneme domain.

**FIG. 6.**
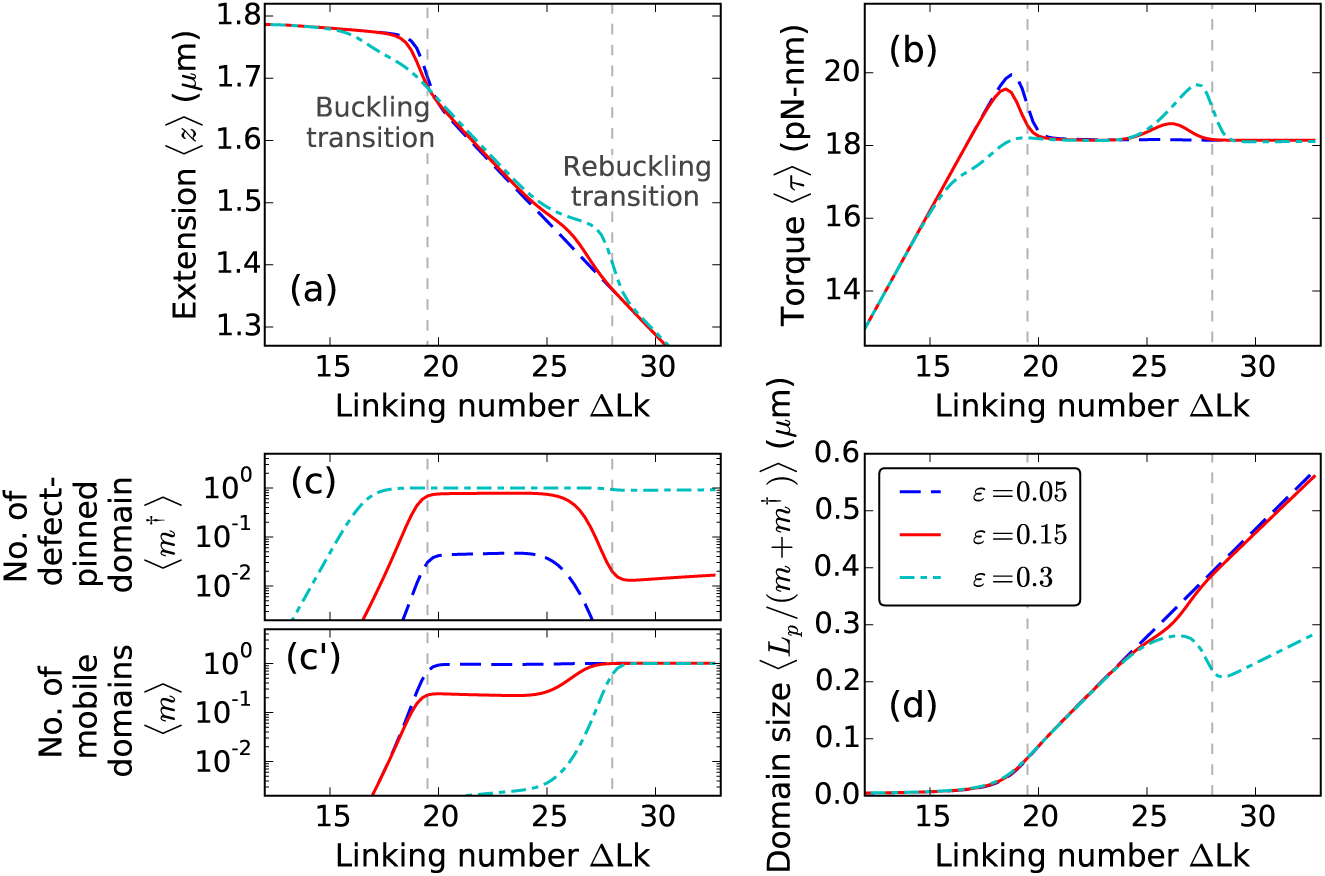
Supercoiling 2*μ*m DNA (≈ 6 kb) with a defect of size *ε* = 0.05 (blue dashed lines), 0.15 (red solid lines), and 0.3 (cyan dot-dashed lines) located L*=150 nm (≈ 450 bp) from the surface, under 2 pN stretching force and 0.5 M Na^+^. The buckling transition is associated with nucleation of a plectoneme domain, whereas, the rebuckling transition is due to a maximum-size constraint on the defect-pinned plectoneme domain [1]. (a) Extension and (b) Torque versus linking number curves show, respectively, a sharp decrease and an overshoot at the buckling and rebuckling transition points. The magnitudes of torque overshoot and extension jump, associated with the nucleation cost at the transition, decrease with increasing defect size *ε* for the buckling transition; whereas, at the rebuckling point, they increase with increasing size of the defect, (c) Equilibrium number of pinned plectoneme domain shows an appearance of the defect-pinned plectoneme at the buckling point, however, probability of nucleating the defect-pinned domain is vanishingly small for *small defects* (*ε* < 0.1). Near the rebuckling point, the defect-pinned domain is stable only for *large defects* (*ε* > 0.25). (c’) Equilibrium number of mobile plectoneme domains shows that a mobile domain is favored at the buckling point only when the defect is small; for larger defects, a mobile domain does not appear before the rebuckling point. This suggests that the rebuckling transition does not occur for small defects, (d) Average size of a plectoneme domain shows an increase after the buckling point. Rebuckling transition occurs when the size of the defect-pinned domain is 2 *L*\* or 0.3 *μ*m. Near the rebuckling point, for larger defects, the average size of a domain shows an abrupt decrease due to nucleation of a mobile plectoneme domain. The vertical dashed lines correspond to buckling and rebuckling transitions [see Figs. 7 and 8], however, note that the critical linking numbers for the transitions are defect size dependent [Fig. 10].

**FIG. 7.**
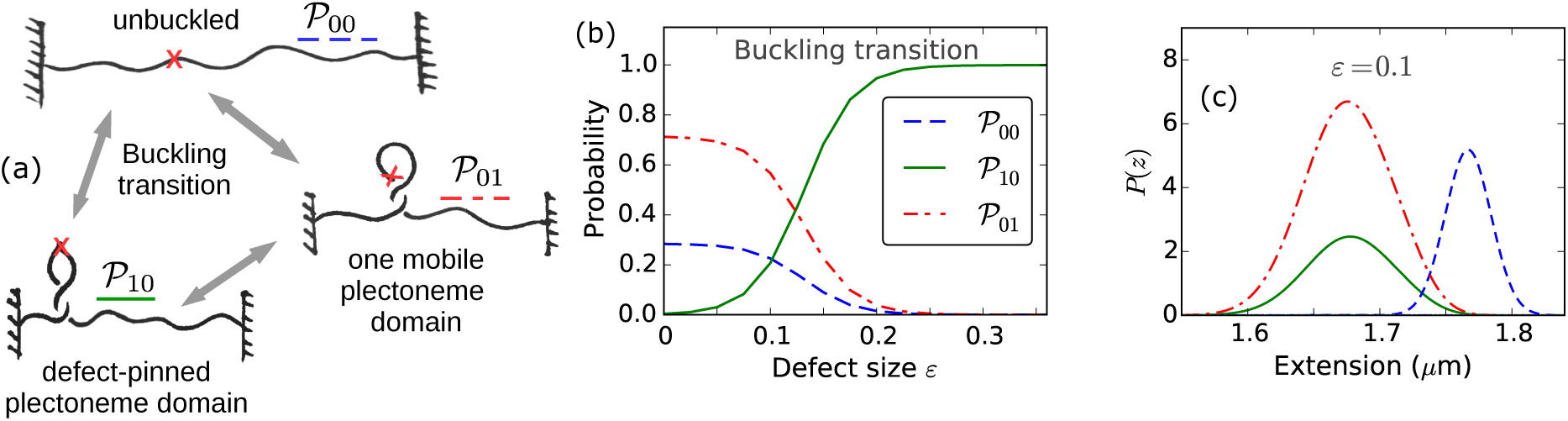
Buckling transition for various defect sizes. (a) Schematic of the three possible states at the buckling transition: unbuckled state (𝒫_00_, blue dashed line), defect-pinned plectoneme (𝒫_10_, green solid line), and mobile plectoneme domain (𝒫_01_, red dot-dashed line). (b) Total probability of the three states at the buckling transition (∆Lk=19.5 under f = 2 pN and 0.5 M Na^+^, see Fig. 6) as a function of the defect size *ε.* For larger defects *(ε* > 0.1), the defect-pinned domain (𝒫_10_) is the favored post-buckling state, because of the lower bending energy of a kinked end loop associated with 𝒫_10_. While, for *small defects* (*ε* < 0.1) the bending energy saved from a kinked end loop is lower than the loss of diffusion entropy of the pinned state (𝒫_10_), which makes the mobile domain ((𝒫_01_) the favored post-buckling state. Note the relatively higher probability of the unbuckled state for smaller defects. This is due to a shift of the buckling point towards lower linking numbers with increasing defect sizes (Fig. 10). (c) Probability density of DNA extension at the buckling transition shows the typical bimodal character observed for defect-free DNA (Fig. 2), however, the defect size controls the states populating the lower-extension mode of the distribution. This also suggests that measurement of the extension alone is insufficient to distinguish between the states involved at the buckling transition.

**FIG. 8.**
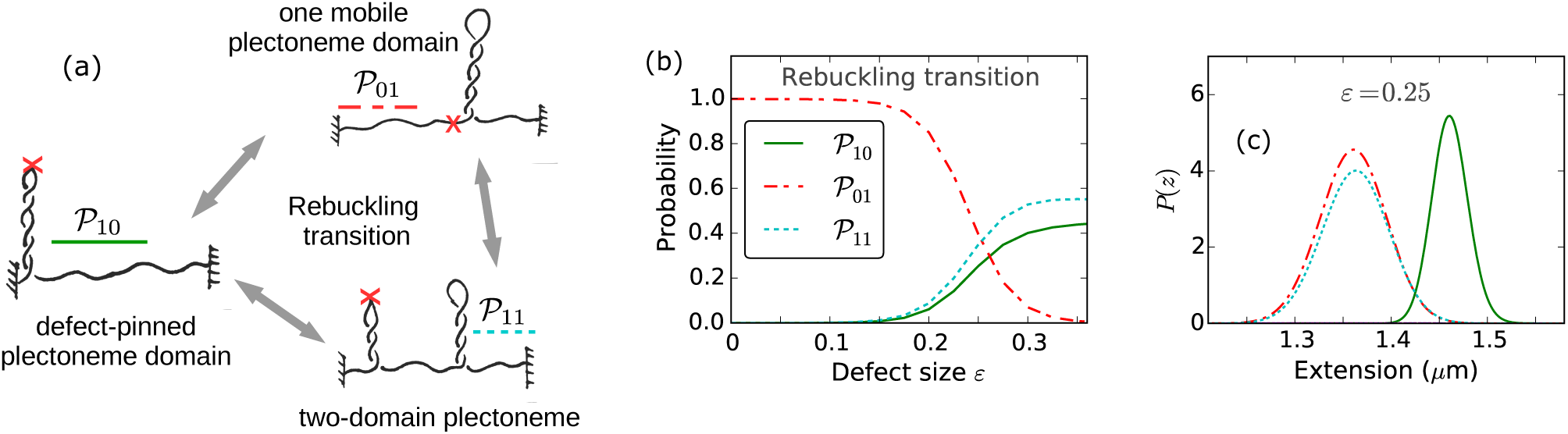
Rebuckling transition for various defect sizes. (a) Schematic of the three possible states at the rebuckling transition: the critically-big defect-pinned plectoneme domain (𝒫_10_, green solid lines), one mobile plectoneme domain (𝒫_01_, red dot-dashed lines), and two-domain plectoneme containing one defect-pinned and one mobile domains (𝒫_11_, cyan dotted lines). (b) Total probability of the three states at the rebuckling point (∆Lk=28 under *f* = 2 pN and 0.5 M Na^+^, see Fig. 6) as a function of the defect size *ε.* For *small defects* (0 < *ε* < 0.1), the post-buckling state is the mobile domain (𝒫_01_) and not the defect-pinned domain (𝒫_10_) (see Fig. 7). As a result, rebuckling is not observed for small defects and the 𝒫_01_ state continues to increase in size after buckling, same as the case for a defect-free DNA (Fig. 2). For *intermediate defects* (0.1 < *ε* < 0.25), the defect is large enough to bias nucleation of the defect-pinned domain (𝒫_10_) at the buckling transition (Fig. 7); however, at the rebuckling point, one mobile domain (𝒫_01_) is more stable than the two-domain state ((𝒫_11_). For *large defects (ε* > 0.25), the defect-pinned domain (𝒫_10_) is highly stable, resulting in nucleation of a new mobile domain at the rebuckling point; this makes the two domain state (𝒫_11_) favored after the rebuckling transition. Note the higher probability of 𝒫_10_ for larger defects, which is due to a shift of the rebuckling transition to higher linking numbers for larger defect sizes (Fig. 10). (c) Probability density of extension for *ε* = 0.25 at the rebuckling point. The bimodal extension profile is due to the finite nucleation energy associated with a teardrop loop of a mobile domain. The state populating the lower-extension mode of the distribution depends on the size of the defect, such that large and intermediate defects favor 𝒫_11_ and 𝒫_01_ states, respectively. Small defects show a unimodal extension profile after buckling transition, and do not exhibit rebuckling.

**FIG. 9.**
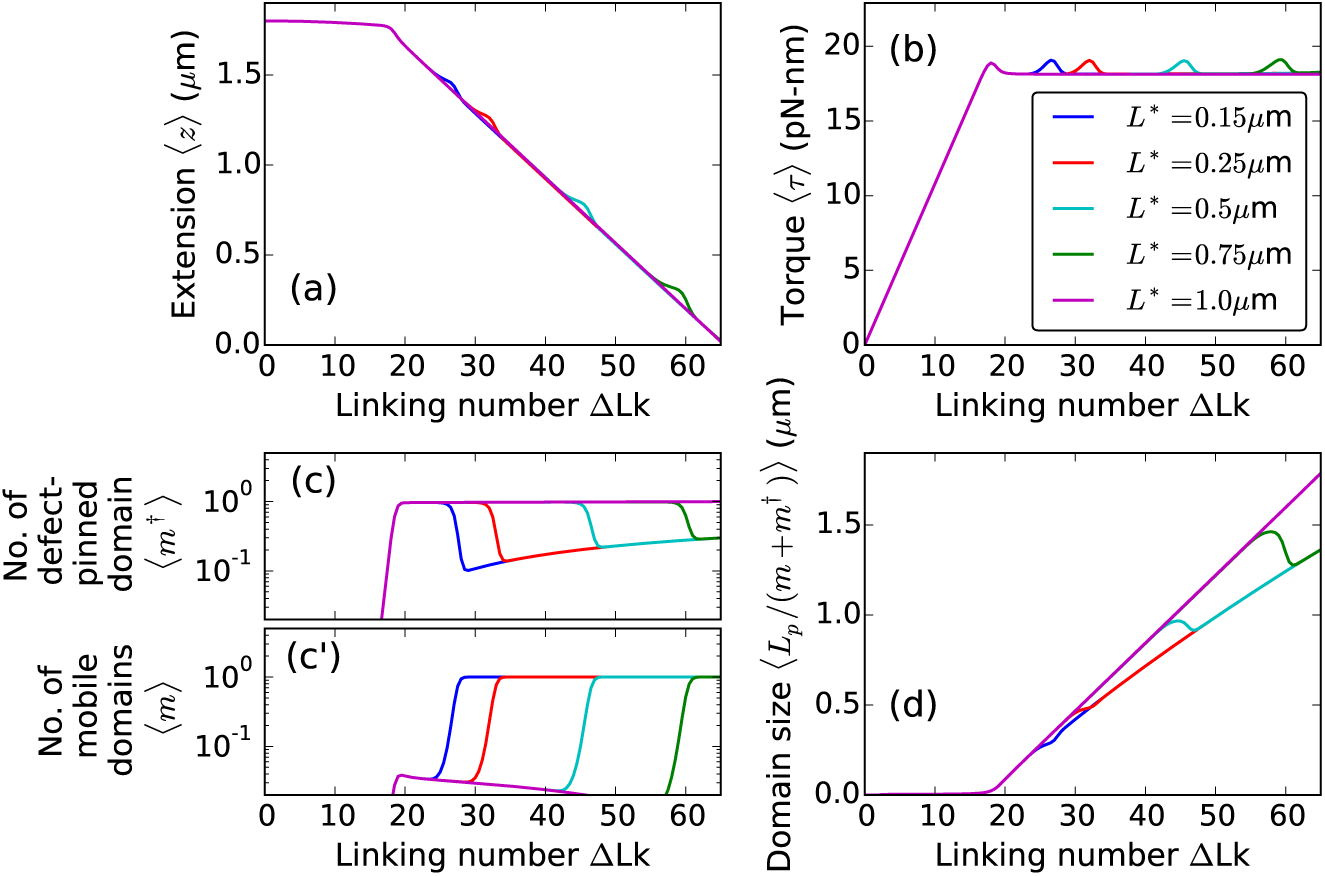
Displacement of the defect site. Twist response of 2 *μ*m DNA under 2 pN force at 0.5 M Na^+^ with an intermediate defect (*ε* = 0.2) located *L\** = 0.15 (blue), 0.25 (red), 0.5 (cyan), 0.75 (green), and 1 *μ*m (magenta) from one of the DNA ends. The location of the defect site controls the critical size of the pinned plectoneme domain (2 *L*\*) nucleated at the buckling point. This results in an increase of the critical linking number for the rebuckling transition for defects located farther away from the end, seen as a shift in extension and torque bumps. For a centrally-located defect (*L*\* = 1 *μ*m, magenta lines) the rebuckling transition does not occur because the critical size of the defect-pinned domain is equal to the total size of the DNA. Note that unpinning of the defect-pinned domain occurs at the rebuckling transition, as expected for intermediate defects.

**FIG. 10.**
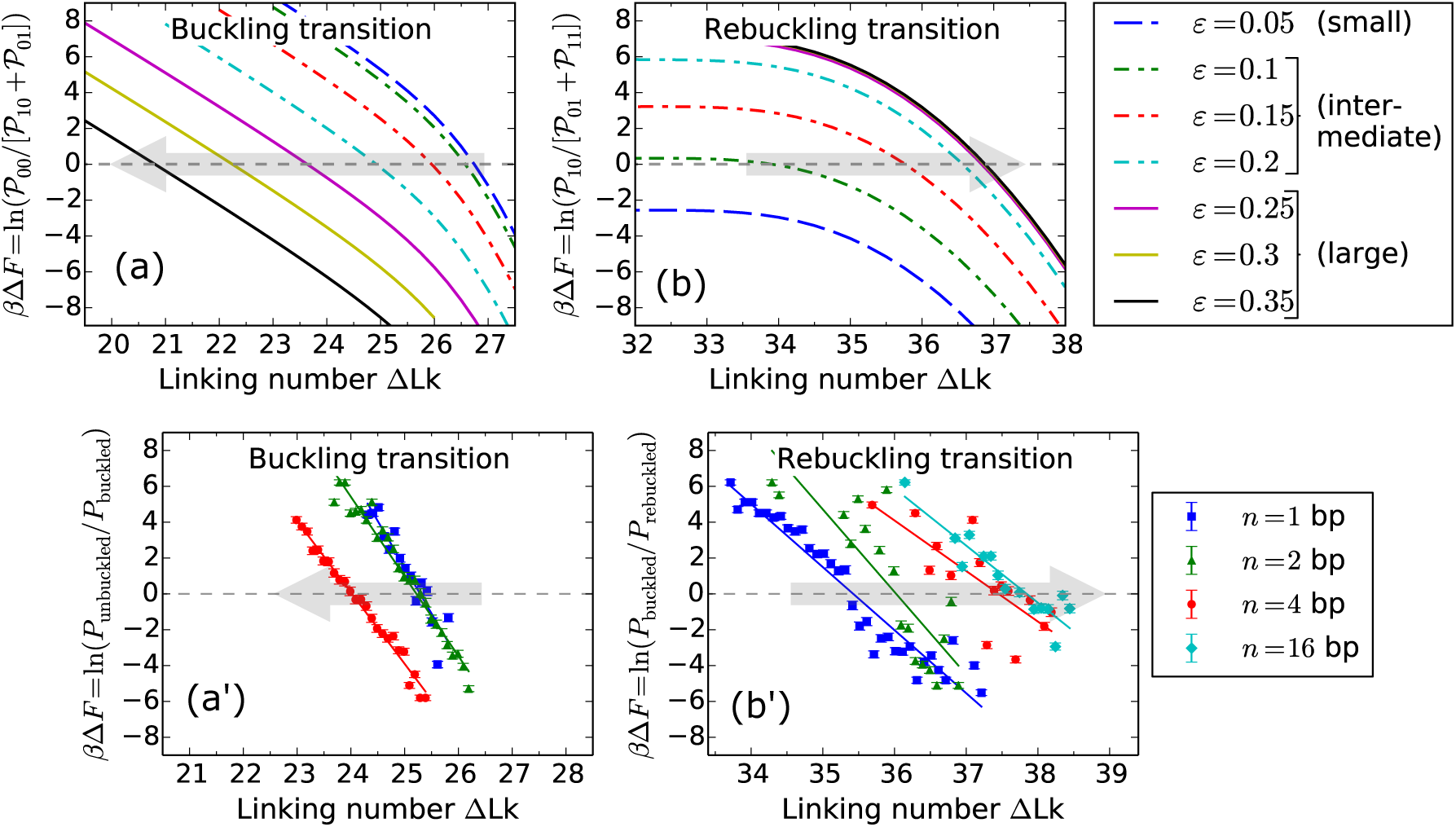
Comparison of theoretical and experimental shifts in the critical linking numbers associated with buckling and rebuckling transitions as a function of the defect size for *f* = 3.6 pN. The size of the defect is defined theoretically via the parameter ε, and experimentally, as *n*, the number of adjacent base pair mismatches on the DNA [1]. Theoretically, the defects are categorized into small, intermediate, and large (Fig. 8) depending on the numerical value of *ε*, as shown in the figure legend. The free energy difference, *∆F* between the lower and higher extension states at the buckling or rebuckling transitions is obtained from the logarithm of the ratio of area-under-curve of extension histograms (as shown on the y-axis labels). For a specified defect size, the critical linking number corresponds to *∆F* = 0. The gray shaded arrows show the direction of increasing defect sizes. (a)Buckling transition: Theoretical plot shows a decrease in the critical linking number with an increase in *ε* for intermediate and large defects, whereas, the buckling point does not shift for small defects. Large and intermediate defects nucleate a defect-pinned plectoneme domain that has a lower nucleation energy, this causes a decrease in the critical linking number. Small defects nucleate a mobile domain at the buckling transition, as a result the critical linking number is independent of the defect size. (a’) A similar shift of the critical buckling point to lower linking numbers with increasing *n* is observed experimentally (see Fig. S4(b) of Ref. [1]), where the solid lines show the best-fit linear regression for various *n.* (b) Rebuckling transition: Theoretical plot showing the change in the critical linking number as a function of *ε.* Small defects do not show rebuckling transition: the blue dashed line does not intersect *∆F* = 0. Intermediate defects show an increase in the associated critical linking number with increasing *ε.* Rebuckling for intermediate defects involves a decrease (increase) in the probability of the defect-pinned domain (one mobile domain), and consequently, a higher stability of the defect-pinned domain delays the rebuckling transition. Large defects show rebuckling but not a shift in the rebuckling point with *ε.* For large defects, a mobile domain is added to the defect-pinned domain at the rebuckling transition, making the rebuckling critical linking number independent of *ε.* (b’) Experimental plot of *∆F* near the rebuckling transition for various *n* (solid curves are the best-fit straight lines) agrees with the theoretical trend of the shift in the rebuckling critical linking number.

#### 1. Buckling transition

At the buckling transition, the plectoneme state becomes energetically favored because of its writhe linking number contribution that decreases DNA-twist energy and torque.

##### a. Coexistence of multiple states

The plectoneme domain nucleated at the buckling transition can be defect-pinned (𝒫_10_), which has a nucleation energy that decreases with increasing defect size, but the spatially-pinned nature of the domain costs diffusion entropy [Fig. 7(a)]. On the other hand, the nucleated plectoneme do main can be a mobile one (𝒫_01_), which has a fixed nucleation cost associated with a teardrop loop and possesses extra stabilization from diffusion entropy. Figure 7(b) shows the probability of the three states: 𝒫_00_, 𝒫_01_, and 𝒫_10_, near the buckling transition as a function of the defect size. The defect-pinned domain (𝒫_10_) is the favored post-buckling state for larger defects (*ε* > 0.1), whereas a mobile domain (Poi) is the preferred post-buckling state for smaller defect sizes (*ε* < 0.1).

##### b. Absence of defect-pinning for small defects

For small defects (0 < *ε* < 0.1), the loss of diffusion entropy for a defect-pinned domain is higher than the bending energy saved from a kinked end loop. This leads to relative stabilization of the mobile domain for small defect sizes. The probability of the mobile plectoneme domain decreases near the buckling transition with increasing defect size, and becomes negligible for larger defects [Figs. 6(c) and 7(b)]. The probability of the defect-pinned plectoneme domain increases with the defect size, and the pinned domain becomes the most probable post-buckling state for larger defects. The total probability of an end loop or a plectoneme domain after the buckling transition is unity; the defect size controls the relative stability of the two possible types of end loops: mobile and kinked or defect-pinned, thus controlling the post-buckled state [Fig. 7(b)].

##### c. Lower critical linking number for larger defects

Buckling occurs at a lower linking number for higher values of *ε*. Figure 6(c) shows an increase in the probability of the defect-pinned plectoneme domain at the buckling point. The nucleation cost of a defect-pinned plectoneme domain decreases with increasing defect size, due to the lower bending energy associated with a kinked end loop of a larger defect [Eq. (8)]. This results in a decrease of the critical linking number for larger defects, as well as a smaller extension change and torque overshoot at the buckling transition [Fig. 6(a)-(b)].

However, small defects do not show a shift in the buckling transition, because the nucleation cost of the most probable post-buckling state (𝒫_01_) is independent of the defect size.

##### d. Plectoneme contribution to the nucleated domain

The average size of a plectoneme domain increases with increasing linking number due to nucleation and consequent growth of a plectoneme domain. Note that although the critical buckling point varies with the defect size, the increase in plectoneme domain size near the buckling point does not depend on the defect [Fig. 6(d)]. DNA length contribution to a nucleated domain from plectoneme superhelical turns depends only on the linking number. As a result, the critical linking number at a transition determines the plectoneme contribution to the total size of the nucleated buckled domain, such that the contribution is larger for a higher critical linking number, i.e., a smaller defect. This also means that the extension change at the buckling transition is larger for smaller defects [Fig. 6].

For small defects *(ε* < 0.1), the domain size increases after nucleation of the mobile end loop. However, for larger defects, the defect-facilitated kinked end loop is highly stable and becomes probable before plectoneme superhelices are favored [Fig. 6(a)-(c)]. As a result, for larger defects, there is a linking number interval at the buckling transition where the post-buckled state is just the kinked end loop with minimal superhelical turns. This interval shrinks as the defect size decreases. Supercoiling experiments using a 20 bp DNA hairpin as a defect have observed such a linking number interval [6].

### 2. Rebuckling transition

In Fig. 6, the defect is located 450 bp away from one of the ends of the 6 kb DNA molecule. As mentioned previously (Sec. III A 2), the position of the defect imposes a critical size of 900 bp or 0.3 µm on the defect-pinned plectoneme domain. This is directly related to the fact that a defect-pinned domain must have the defect site at its tip, where an energy-saving kinked end loop is placed [Fig. 1(b)]. When a defect-pinned domain nucleated at buckling becomes critical in size, nucleation of a mobile plectoneme domain is required to store additional linking number as writhe, resulting in an increased probability of a mobile domain at the rebuckling transition [Fig. 6(c^′^)].

The nucleation of a mobile plectoneme domain may or may not be accompanied with a reduction in the equilibrium probability of the defect-pinned domain (𝒫_10_). This leads to two possible post-rebuckling states: one mobile plectoneme state (𝒫_01_), and the two-domain plectoneme state (𝒫_11_) containing one mobile domain and a defect-pinned domain. Figure 8(a) shows a sketch of the three coexisting states at the rebuckling transition. The corresponding probability of occupancy of these states at the rebuckling point are also plotted for various defect sizes [Fig. 8(b)].

#### a. Nucleation cost at rebuckling

The nucleation energy of the post-rebuckling state depends on the relative stability of the defect-kinked end loop with respect to a teardrop loop, such that the nucleation cost of the mobile plectoneme state (𝒫_01_) increases with the defect size. Note that the total nucleation cost of the 𝒫_01_ state at the rebuckling point is associated with, first, the energy cost of a teardrop loop which is defect-size independent, and second, the cost of *unpinning* a defect-pinned domain (i.e., a decrease in the equilibrium probability of the pinned domain) which increases for larger defects. While, the nucleation cost for the two-domain state (𝒫_11_) depends only on the energy of the teardrop loop of the added mobile domain and does not change with the defect size. The discontinuity in extension and overshoot in torque, seen at the rebuckling point [Fig. 6(a)-(b)], is due to the finite nucleation cost associated with the post-rebuckling state.

#### b. Absence of rebuckling for small defects

Small defects (0 < ε < 0.1) do not exhibit rebuckling transition because the defect-pinned domain is not the most probable post-buckling state [Fig. 6(a)-(c)]. For small defects, a mobile plectoneme (𝒫_01_) is nucleated at the buckling transition which increases in size with increasing linking number in the buckled state. Since a mobile plectoneme domain (𝒫_01_) is the most probable buckled state for small defects, the probability corresponding to 𝒫_01_ is the highest for small defect sizes in Fig. 8(b). For small defects, the defect-pinned domain is the second most probable post-buckling state, and the probability goes to zero when the total plectoneme size is larger than its critical size [Fig. 8(c)].

#### c. Unpinning of the defect-pinned domain for intermediate defects

For intermediate defects (0.1 < *ε* < 0.25), probability of the defect-pinned domain decreases at rebuckling. This means that the critically-big defect-pinned domain is “unpinned” or transformed into a mobile domain by displacing it along the DNA contour, thereby replacing the kinked end loop with a teardrop end loop. Hence, for intermediate defects, the pre-rebuckling state is the defect-pinned domain (𝒫_10_) and the post-rebuckling state is a mobile plectoneme domain (𝒫_01_) [Fig. 8(b)].

#### d. Two-domain plectoneme for large defects

For large defect sizes (*ε* > 0.25), higher stability of the defect-pinned domain resists decrease in its equilibrium probability and results in addition of a mobile plectoneme domain [Fig. 6(c′)]. Thus, for large defects, the favored post-rebuckling state is the two-domain plectoneme state (𝒫_11_); whereas, the pre-rebuckling state is the defect-pinned domain (𝒫_10_), which is essential for rebuckling to occur [Fig. 8].

#### e. Critical linking number increase with increasing defect size

For intermediate defects, the rebuckling transition occurs at a higher linking number for larger defect sizes, which is related to the increased stability of the defect-pinned domain [Fig. 6]. This is because the nucleation cost at rebuckling increases with increasing defect size resulting in an increase in the associated critical linking number. However, for large defects, the nucleation cost does not depend on the defect size resulting in a critical linking number that does not change with the defect size. The shift in the critical linking number with defect size is also seen experimentally [1], and is quantitatively analyzed later in this article [Fig. 10].

#### f. Absence of rebuckling at lower stretching forces

At lower forces, the nucleation cost of a plectoneme domain decreases, resulting in an increased tendency to proliferate multiple domains. The energy difference between a mobile and a kinked end loop also decreases with decreasing force, leading to a coexistence of the defect-pinned and mobile plectoneme domains in the post-buckling state. For intermediate defects, below ≈ 1 pN the mobile domain is the most probable post-buckling state, whereas, the defect-pinned domain is the second most probable. Hence, intermediate defects do not show rebuckling transition when the stretching force is less than ≈ 1 pN.

#### g. Effect of lowering the salt concentration

Lower salts promote proliferation of multiple plectoneme domains due to decreased stability of plectoneme super-helices that contain close proximity of DNA segments (Fig. 4). This causes an increase in the probability of the two-domain state at the rebuckling transition as the ionic strength of the solution is decreased. Hence, at salt concentrations ≈ 0.1 M Na+, intermediate defects preferentially nucleate the two-domain state (𝒫_11_) at the rebuckling transition.

#### h. Displacing the defect shifts the rebuckling point

For a given force and length of the molecule, the critical linking number associated with rebuckling depends on the location of the defect. A defect located farther away from the closest end of the DNA (i.e., larger *L*)* increases the critical size of the defect-pinned domain, resulting in an increase in the rebuckling critical linking number (Fig. 9). Note that for a defect located at the middle of the DNA, the defect-pinned domain is critically big only in the purely plectoneme state (no unbuckled DNA or zero extension), and the rebuckling transition does not occur, as has also been shown experimentally [1].

## 3. Experimental detection of three-state coexistence

The extension distributions at the buckling and rebuckling transitions appear bimodal, even when there are three coexisting states [Figs. 7(c) and 8(c)]. The lower-extension peak at the transition is the sum of contributions from the two possible post-transition states (𝒫_01_ and 𝒫_10_ for buckling, and 𝒫_01_ and 𝒫_11_ for rebuckling). This means that experimentally measuring the extension profile at the buckling or rebuckling transitions, as done in magnetic tweezer experiments [1], does not inform on the identity of the post-transition state.

Measurements of the lifetime of the lower and higher extension modes are also unlikely to shed light on the possibility of multiple states contributing to the lower-extension peak. The lifetimes of the higher and lower extension states are simple exponential distributions. In case of a three-state transition where one of the states is *hidden* (i.e., the transition out of one state is not the same as the transition in to the other and vice-versa), the lifetimes follow a Gamma distribution (polynomial increase for small times and exponential decay for long times). However, if two of the three states are grouped together, like the predicted scenario, the transition out of one state is the same as the transition in to the other and vice-versa, resulting in an exponential distribution of the lifetimes. Such grouping of the two states in a three-state transition simply blinds the observer to ~ 1/3 of the transition events and the overall kinetics appears to be that of a two-state transition.

However, DNA-visualization experiments, where the DNA backbone is labeled with a fluorescent dye [5, 16], may be able to distinguish between a defect-pinned domain, a mobile domain, and a two-domain plectoneme state. Our model predicts the most probable state for a given linking number and force, but does not inform on the kinetic pathways at the transition. Precise control of the DNA linking number in visualization experiments may also be able to report on kinetically-favored states at buckling and rebuckling transitions.

## C. Comparison with Experiments

In this section we compare our results with magnetic tweezer experiments on supercoiled DNA with a base-unpaired region [1]. We obtain a quantitative relation between the number of adjacent base mismatches n, and the theoretical defect size *ε.* Experiments and theory show good agreement for various thermodynamic trends.

### 1. Critical linking number shift

#### a. Buckling transition

The critical linking number associated with buckling shows a general decreasing trend with increasing defect size, because of the lower nucleation cost for larger defects [gray arrows in Fig. 10(a)- (a’)]. We define the critical linking number as the point where the higher and lower extension peaks have equal weights (i.e., *∆F* = 0 in Fig. 10). Experiments show a similar shift in the buckling point to a lower linking number as the number of adjacent base mismatches *n* is increased [1] [Fig. 10(a’)]. For small defects, the nucleation cost at buckling transition is less sensitive to the defect size because of the lower probability of the defect-pinned domain [Fig. 7(b)], resulting in a very small to no shift in the buckling point [Fig. 10(a)].

#### b. Rebuckling transition

The rebuckling point shifts to higher linking numbers with increasing defect size for intermediate defects. As previously mentioned, rebuckling does not occur for small defects [In Fig. 10(b), the blue dashed curve corresponding to a small defect does not intersect *∆F =* 0]. For large defects, the critical linking number for rebuckling transition does not depend on the defect size. The nucleation cost of the post-transition state (two-domain plectoneme, 𝒫_11_) for large defects is associated with a mobile end loop and does not depend on the defect size. Experiments show the expected trend for the rebuckling point shift [Fig. 10(b’)]. Note that *n =* 4 and 16 bp cases show a very small shift in the rebuckling point, the expected behavior for large defects.

We compare theoretical results for 0.5 M Na^+^ with experimental observations at 1 M NaCl [1] [Figs. 10 and 11]. The increased abruptness of the buckling and rebuckling transitions at higher salt concentrations, related to the higher stability of plectoneme superhelices, makes the high salt scenario suitable for experimental studies. However, at salt concentrations higher than 0.5 M, the Debye-Hückel approximation becomes questionable at best.

**FIG. 11.**
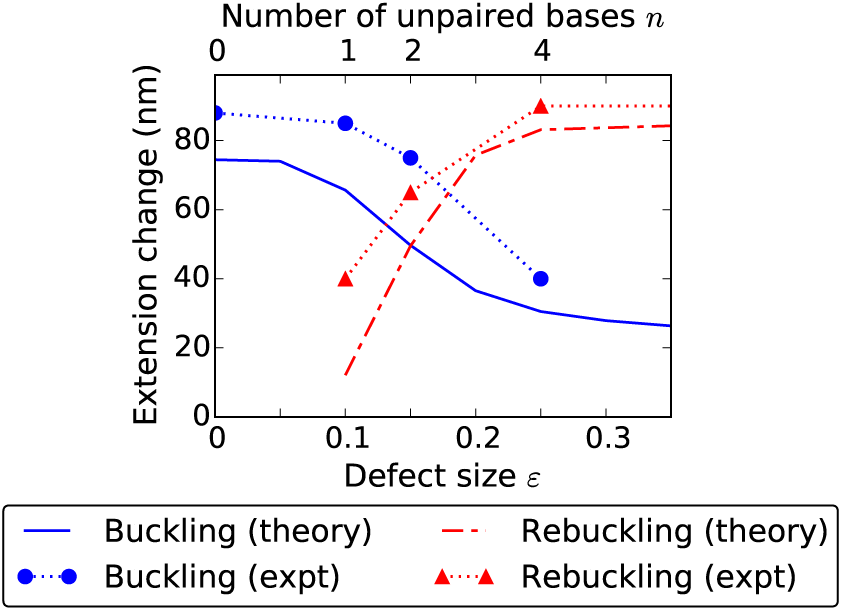
Change in extension at the buckling and rebuckling transitions as a function of the defect size for 3.6 pN force. The extension change at a transition depends on the size of the nucleated domain, which has contributions from the end loop and the plectoneme superhelical state. In case of buckling (blue solid line), the critical linking number decreases with increasing defect size causing a decrease in the plectoneme contribution as well as the size of the end loop, reducing the extension change at the transition. However, for small defects, nucleation of a mobile domain causes a saturation in the extension change for buckling transition and an absence of rebuckling transition, hence no associated extension change (red dot-dashed line). For intermediate defects, rebuckling extension change increases with the defect size due to an increase in the associated critical linking number resulting in a higher superhelix contribution. Whereas, for large defects, the extension change is constant because of a fixed superhelix contribution corresponding to a fixed critical linking number (Fig. 10). Experimental data for the extension change (see Fig. 2(c) in Ref. [1]) as function of the number of adjacent unpaired bases *n* (top x-axis) compares well with theory. The experimental error bars, omitted in the plot, are smaller than the size of the point markers.

### 2. Relation between ε and n

The experimental (n) and theoretical (*ε*) defect sizes are expected to have a direct monotonic relationship for *n* ≥ 1, and *ε* = 0 for *n =* 0. We look for a simple linear variation for *n* ≥ 1: *ε* = *a + bn*, ignoring higher order terms that only contribute for very large defect sizes.

The fact that rebuckling is observed experimentally for *n =* 1 suggests that it is not a small defect. The rebuckling point shifts in the experimental plot with increasing defect size [Fig. 10(b’)]; this indicates that at least *n =* 1 and 2 must be intermediate defects. Comparing the critical linking number shifts we see that an increase in *n* by 1 roughly corresponds to an increase of 0.05 in *ε*, implying *b* ≈ 0.05.

The smallest defect sizes corresponding to rebuckling transition are *ε* ≈0.1 and *n =* 1; this suggests *a+b* ≈ 0.1. Hence, we find *n =* 1,2, and 4 respectively correspond to *ε* = 0.1,0.15, and 0.25.

The bending energy saved by increasing the defect size by Δ*ε* is 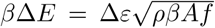 [Eq. (8)]. This suggests that increasing the size of a DNA lesion by one base pair (Δ*ε* = 0.05) saves ≈ 0.7 *k*_B_*T* of bending energy by allowing a sharper kink at the lesion site under physiological forces (1 pN). Our estimation is based on intermediate defects, and may be an underestimation for larger defect sizes because, first, we assumed a linear relationship between *n* and *ε*; and second, we ignored any defect-induced perturbation in DNA twist.

One may ask, given that a larger defect stabilizes the kinked end loop, when is it favorable to break the intact base pairs adjacent to a base-unpaired region, i.e., save bending energy by increasing the defect size at the cost of base-pairing energy? The energy saved for an increase in the defect size by 1 bp (Δ*ε* = 0.05) when equated with the average base-pairing energy ≈ 2.5 *k-gT* [47], we find a critical force ≈ 12.5 pN which compares with DNA unzipping force ≈ 12 pN [48]. This means that the base-unpaired region of a supercoiled DNA will show an equilibrium increase in size when stretched under forces larger than ≈ 12.5 pN. This value, based on our estimation of the defect size, provides a reference point. Experimental significance of this force is not clear because we ignore transitions in the double-helical structure of DNA, which are known to occur at torques ≳ pN-nm, which corresponds to buckling torque under ≳ 6 pN stretching force [49].

### 3. Abruptness of the transition

The extension change at a transition is a measure of the abruptness of the transition. Transitions with more resolved peaks in the bimodal extension profile are more abrupt.

#### a. Buckling transition

At the buckling transition, the total size of the nucleated domain (lower extension state) has contributions from the end loop (kinked or mobile) and the plectoneme superhelical structure. Larger nucleation cost increases the critical buckling linking number which increases the amount of plectonemic turns in the nucleated domain. As the defect size gets bigger (intermediate and large defects), the nucleation cost decreases, which reduces the plectonemic contribution to the nucleated domain, resulting in a decrease of the extension change [Fig. 11]. Note that the size of a kinked end loop at 3.6 pN is 30 – 20 nm for defect sizes 0 < *ε* < 0.35 [Fig. 11, and Eq. (9)], and it goes to zero as *ε*→1.

Hence, the steep decrease in the extension change at the buckling transition for intermediate defects is predominantly due to a decrease in the plectoneme contribution to the nucleated domain [Fig. 11]. A large defect nucleates a kinked end loop with minimal plectoneme. This makes the extension change depend solely on the size of the kinked end loop, and the small change in the kinked end loop length with increasing defect size results in the shallow decrease in extension change for larger defects [Fig. 11]. Small defects nucleate a mobile domain, resulting in an extension change that does not depend on the defect size [Fig. 11].

#### b. Rebuckling transition

Extension change at the rebuckling transition increases with increasing defect size for intermediate defects. Rebuckling occurs at a higher linking number for larger defects due to increased nucleation cost, resulting in an increased plectoneme contribution to the nucleated domain. This means that at the rebuckling point, total plectoneme length in one mobile plectoneme domain (𝒫_01_, the favored post-rebuckling state for intermediate defects) is larger than that for the defect-pinned plectoneme domain (𝒫_10_, the pre-rebuckling state). The plectoneme contribution to the post-rebuckling state also increases with the defect size, resulting in an increase of the rebuckling extension change [Fig. 11].

Large defects nucleate the two-domain plectoneme at the rebuckling point, which has a defect-independent nucleation cost, and as a result the plectoneme contribution to the nucleated domain does not depend on the defect size. This produces a fixed extension change for rebuckling in large defects [Fig. 11]. Small defects do not show rebuckling, hence there is no extension change associated with small defects at rebuckling.

### 4 Detection of rebuckling signal in force-salt landscape

Experimental detection of the rebuckling transition relies on the resolution of the corresponding bimodal extension distribution [1]. As the force and ionic strength are lowered, the rebuckling signal diminishes due to overlapping peaks in the extension profile that gives the distribution an overall unimodal character. Consequently, rebuckling is experimentally observed mainly in the high salt and high force regime [1].

### Pearson chi-squared test

We implement a chi-squared analysis that compares the theoretical extension distribution to a single Gaussian that best fits the distribution. We fit the theoretical total-extension histograms [P(*z*), Eq. (13)] near the rebuckling point to a Gaussian distribution, where the mean and the variance are obtained via least-squared method. We then use a Pearson chi-squared test [50] to find the p-value of the fit; the p-values corresponding to various salts and forces at the rebuckling transition are plotted in Fig. 12.

**FIG. 12.**
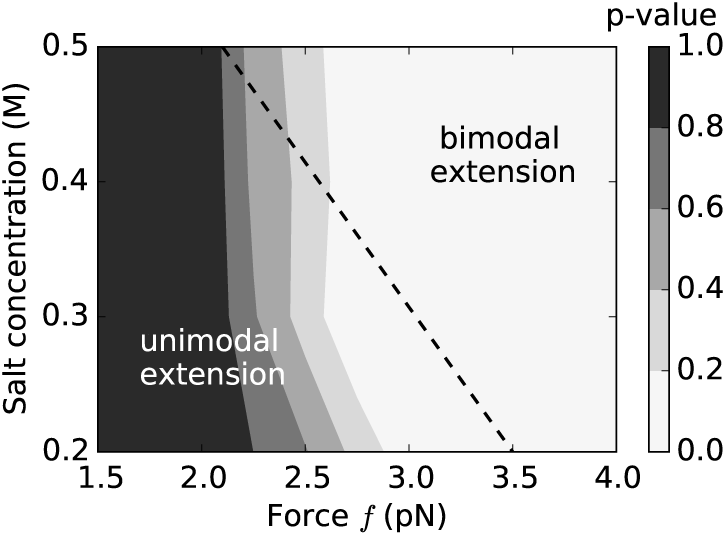
Contour plot of p-values near rebuckling transition (*ε* = 0.15) for various points on the force-salt landscape. We fit the theoretical extension distributions near the rebuckling point to a single Gaussian distribution, and using the chi-squared test calculate the p-value, which serves as a goodness-of-fit statistic. Lower p-values (lighter shade) indicate that the extension histograms near the rebuckling point are characteristically bimodal and are not well fitted by single Gaussian distributions; whereas, higher p-values (darker shade) indicate the rebuckling extension histograms are well approximated as Gaussian distributions. Since the experimental signal associated with the rebuckling transition is the bimodal character of DNA extension, we find that the rebuckling transition is more likely to be observed experimentally when the p-value is low, i.e., higher forces and higher salts. The dotted black line shows the experimentally observed boundary for the appearance of the rebuckling signal associated with a defect containing two adjacent base-pair mismatches *(n* = 2 bp) (see Fig. 3 of Ref. [1]). The bimodal rebuckling signal was reliably observed in experiments for salts and forces on the right-hand side of the dotted line.

Low p-values indicate that our null hypothesis of the chi-squared test: *extension histograms are Gaussian*, is less likely, which arises from the distinctly bimodal nature of the extension profile. As a result, low p-values correspond to the regions in the force-salt landscape where rebuckling produces an extension profile with two resolvable peaks, and is more likely to be observed experimentally. On the other hand, higher p-values suggest that the total-extension profile is well fit by a single Gaussian distribution (unimodal), and experimental detection of rebuckling is less probable.

The dotted line in Fig. 12 shows the experimentally-observed boundary between the disappearance of the rebuckling signal on the left-hand side of the dotted line, i.e., at lower forces and salts; and the appearance of the signal on the right-hand side of the dotted line, i.e., for higher forces and salt concentrations [1]. The theoretical picture correlates well with the observed statistics of the rebuckling signal, where bimodal extension profile is expected for higher salts and higher forces (low p-value) [lighter shade in Fig. 12]. We note that the experimental boundary [dotted line in Fig. 12] is associated with a statistical disappearance of the experimental rebuckling signal [1]; and theory suggests that this disappearance is not due to an absence of the rebuckling transition, but thermal fluctuations overshadowing the experimental signal at lower forces and salts.

## IV. CONCLUSIONS

### A. Defect-Free DNA

Supercoiled defect-free DNA shows a linear torque buildup and small extension change upon small linking number change in the DNA (Fig. 2). Larger torques for higher linking numbers drive coexistence of a buckled plectoneme state that is favored due to its substantial writhe linking number contribution [Fig. 1(a)]. DNA torque is nearly constant in the plectoneme-coexistence state due to plectonemic writhe. Torque, however, increases in the purely-plectoneme state due to an increase in DNA twist with increasing linking number (Fig. 3).

#### a. Abrupt plectoneme-buckling transition

The buckling transition marks the onset of a plectoneme-coexistence state. Nucleation of a plectoneme domain is abrupt due to the finite-energy cost associated with a plectoneme end loop. The nucleation cost is also related to the discontinuous change in extension and overshoot in torque at the buckling transition (Fig. 2).

#### b. Multiple plectoneme domains in long DNA molecules

Longer DNA molecules have a larger configuration entropy associated with a plectoneme domain that reduces the nucleation cost of the domain. This results in an increase in the equilibrium number of plectoneme domains coexisting in buckled DNA (Fig. 3). However, in the purely-plectoneme state of the DNA, a single plectoneme domain is favored and the post-buckling torque increases (Fig. 3).

#### c. Highly stable plectoneme at higher salts

Increased salt concentration of the solution causes an increased screening of the DNA charge that leads to highly stable plectoneme superhelices. Whereas, at lower salts, DNA loops are favored over plectoneme superhelices (Fig. 4). This reduces the DNA-length contribution of plectoneme superhelices to the nucleated buckled domain at the buckling transition for lower salts, resulting in a lower extension discontinuity or more rounded buckling transition.

The extension distribution is bimodal in the buckled state for low salt concentrations due to proliferation of multiple domains in the plectoneme-coexistence state (Fig. 5), which may be possible to observe in magnetic tweezer experiments.

#### d. Plectoneme tails

A plectoneme tail is a finite-sized structure connecting the plectoneme domain to the unbuckled part of the DNA. The energy cost of a tail region is related to the constraint of a continuous change of DNA curvature from the plectoneme to the unbuckled DNA, and compares with the energy cost of constraining the ends of a DNA loop (≈ 10% of the bending energy of a loop, i.e., ≈ 2 *k*_B_*T* under 2 pN force, see Sec. IVA of Ref. [37]). The plectoneme tail causes a small increase in the nucleation cost of the plectoneme domain, which we have ignored in our model for simplicity.

However, the tail region may induce a preference for the spatial location of the plectoneme domain. By placing the plectoneme domain at one of the ends of the DNA, so that its tail coincides with a DNA-tether point, the energy associated with the tail can be halved. This is because one of the ends of plectoneme is the tether point and the constraint of a continuous DNA curvature is released for that end. Lowering the plectoneme tail energy by ≈ 1 *k*_B_*T* results in ≈ 3-fold increase in the probability of localizing the domain at one of the DNA ends. There is experimental evidence for preferential localization of a plectoneme domain near the DNA end [5].

## B. DNA with a Defect

We analyzed the effect of an immobile point defect, e.g., a short base-unpaired region [1], on the mechanical response of DNA. We hypothesized that a defect allows a kink on the DNA at the defect site, and the location of the defect imposes a critical size on the plectoneme domain that nucleates at the defect site. A plectoneme domain with the tip of its end loop placed at the defect site (i.e., the defect-pinned plectoneme domain), is energetically favored due to a lower-energy kinked end loop. However, the requirement of putting the end loop at the defect site causes the other end of the plectoneme (i.e., the plectoneme tail) to coincide with a DNA tether point for a certain size of the defect-pinned plectoneme domain, which is its critical size. From simple geometry, a critically-big defect-pinned domain cannot store more superhelical turns [Fig. 1(b)].

In previously reported experiments [1], we used one or more adjacent base pair mismatches at a specified location on the DNA as a point defect. Twisting a base-pair mismatched DNA showed rebuckling transition near the point where the defect-pinned plectoneme domain reaches its critical size [Fig. 6].

### a. Defect size

A defect of size *ε* can nucleate akinked end loop that has a bending energy equal to (1-ε) times the bending energy corresponding to a teardrop loop [Eq. 8]. The defect size *ε* is related to *n* the number of adjacent base-pair mismatches on the DNA, such that a larger value of *n* indicates higher *ε.* We categorized the defect size into: small (0 < *ε* < 0.1), intermediate (0.1 < *ε* < 0.25), and large (0.25 < *ε* < 1).

### b. Buckling transition

Twisting the double helix containing a defect buckles when it is energetically favorable to convert twist into writhe. For intermediate and large defects, the buckling transition occurs via nucleation of a kinked end loop, which makes the defect-pinned plectoneme domain favored at the buckling transition (Fig. 7). For small defects, a mobile plectoneme domain is nucleated at the buckling transition, the DNA kink at the defect reduces bending energy but the defect-pinned domain lacks stabilization from diffusion entropy. This leads to a higher net nucleation cost for the defect-pinned domain featuring small defects than a mobile domain.

The critical linking number associated with the buckling transition decreases with increasing defect size for intermediate and large defects. This is because of the lower nucleation cost associated with a more sharply kinked end loop of a larger defect size (Fig. 6). However, the critical linking number does not change for small defects, because a mobile plectoneme domain is nucleated at the buckling point.

### c. Rebuckling transition

As the linking number is increased after the buckling transition, the defect-pinned domain grows in size and becomes critically big. Further increase in DNA linking number leads to nucleation of a mobile plectoneme domain, required to store additional plectonemic-superhelical turns.

For large defects, the high stability of the kinked end loop leads to addition of a mobile domain such that the post-rebuckling state is that of a two-domain plectoneme featuring one mobile and one defect-pinned domains (Fig. 8). For intermediate defects, the defect-pinned domain unpins itself to nucleate a mobile plectoneme domain. Since only one domain is favored due to high stability of plectoneme superhelices (higher salt concentration scenario), the critically-big defect-pinned domain decreases in probability as the mobile domain becomes more probable after the rebuckling point. For small defects, the rebuckling transition does not occur as the defect-pinned domain is not the most probable after buckling transition (Fig. 8).

For intermediate defects, the critical linking number associated with rebuckling transition increases with increasing defect sizes. This is due to an increased stability of the defect-pinned domain for larger defect sizes (Fig. 6). However, for large defects, the critical linking number does not change because the energy difference between the pre- (critically-big defect-pinned domain) and post-rebuckling (two-domain plectoneme) states does not depend on the defect size.

Lowering the salt concentration of the solution results in an increased proliferation of buckled domains. This increases the probability of the two-domain state for lower salts. Consequently, at low salt concentrations (< 0.1 M Na^+^) unpinning of the defect-pinned domain is not favored for intermediate defects.

An unpinned mobile domain may have a preference for placing its tail at one of the DNA ends or the defect site. This is due to the energy saved from an unconstrained plectoneme tail at the DNA ends or the defect site, which is expected to increase the spatial plectoneme density at those regions and may be visible in fluorescent-DNA experiments [5, 16].

### d. Number of unpaired DNA bases n, and defect size ε

We compare theoretical and experimental shifts in the critical linking numbers at buckling and rebuckling transitions for various defect sizes (Fig. 10). Experimentally, the defect size is controlled via *n* the number of adjacent unpaired bases on the DNA [1], which has a direct monotonic variation with the theoretical defect size ε. We find that *n =* 1,2, and 4 roughly correspond to *ε* = 0.1,0.15, and 0.25, respectively (Fig. 10). This indicates that, for intermediate defects, increasing the size of a base-unpaired region by 1 bp saves ≈ 0.7 *k*_B_*T* of bending energy under 1 pN stretching force.

We predict *n =* 1 and 2 to be intermediate defects, i.e., the defect-pinned domain unpins itself at the rebuckling transition, as opposed to nucleating a two-domain plectoneme under higher salt concentrations (≈ 0.5 M Na^+^) [Fig. 8(b)]. The extension histograms at the rebuckling transition are bimodal even though there are three states: both the mobile and the two-domain plectoneme state contribute to the lower-extension mode [Fig. 8(c)]. As a result, measuring the extension, as done in magnetic tweezer experiments [1] does not inform whether the post-rebuckling state is a mobile domain or a two-domain state. However, DNA visualization using fluorescence imaging experiments may elucidate the existence of these states.

We assume that only the bending degree of freedom of DNA (and not the twist degree of freedom) is affected by the presence of the defect. This assumption is a simplifying one. Certain defects, such as the one produced by a base-unpaired region on the DNA [1], may absorb twist at the defect site. A more sophisticated model may consider the defect to have a lower twist modulus than the double helix, which is essential when treating the base-unpaired region as a coexisting state [51]. However, when the number of adjacent unpaired bases is only a few compared to thousands of paired bases, twist absorption at the defect is expected to be a small effect and may be ignored.

### e. Extension change at the transitions

The change in extension at a transition is non-zero because the nucleated plectoneme domain cannot be smaller than the end loop. Moreover, the relative stability of the end loop compared to plectoneme superhelices determine the contribution of the plectoneme-superhelix state to the nucleated domain; such that when superhelices are highly stable, the plectoneme content of the nucleated domain is larger and results in a larger extension discontinuity. The extension change decreases with increasing defect size at the buckling transition because of a decrease in the plectoneme contribution to the nucleated domain as well as a decrease in the size of the kinked end loop for larger defects (Fig. 11). The extension change at the rebuckling transition increases with an increase in the defect size for intermediate defects, and saturates for large defects.

### f. Disappearance of experimental rebuckling signal at lower force and salt

The extension distributions at the rebuckling transition are bimodal with well-resolved peaks at higher salts and forces (Fig. 12). However, for lower salts and forces, the two peaks overlap giving the extension distribution a unimodal character that obscures experimental detection of rebuckling transition. Lower stability of plectonemic superhelices at lower salt concentrations produces more rounded transitions and masks the rebuckling signal [1].

### g. Biological significance of a defect

The parameter *ε* directly controls the amount of bending energy saved when the tip of a plectoneme end loop is placed at the defect location. The biological relevance of a defect is diverse. Adjacent base-pair mismatches on the double-helix backbone can introduce a spatially-pinned defect [1, 5]; alternately, local structural rearrangement of the double helix by a protein, thereby allowing easy local bending of the DNA backbone [30, 31], might also act as an immobile defect site. Double helices with a single-stranded bulge are a common substrate in single molecule study of various DNA-binding proteins [52]; DNA bulges can also be treated as a defect. The defect size parameter ε may be used as a common scale to compare relative perturbations introduced by defects of varied origin.

Spatial pinning of a plectoneme domain by a defect may have relevance in double-helix base-pair repair mechanism in the cell [1]. A common mechanism to locate targets in the cell, such as locating a DNA base-pair mismatch region by the repair machinery, is that of a diffusive search [53–55]. Preferential positioning of the defect site containing mismatched base pairs at the tip of a plectoneme may facilitate easier access to the lesion.

Moreover, DNA kinks are known to stabilize binding of the enzymes associated with the repair process [56].

### h. Defect induced by DNA sequence

Spatial inhomogeneity in a DNA polymer is not only from varied intra-base-pair interactions, but stacking interactions between adjacent base pairs can enhance the inhomogeneity locally for certain positioning sequences (or, nucleosome-positioning sequences) [26–29]. Pinning of a plectoneme domain by certain sequences have recently been demonstrated experimentally [57]. Occurrence of such positioning sequences in a DNA with otherwise random base pairs may be modeled as a spatially-pinned defect. Local stiffness change from one base pair or a weak sequence-induced defect may be expected to be small, i.e., buckling is not necessarily favored at the defect site and rebuckling is not observed. On the other hand, some positioning sequences may generate an intermediate or large defect, thereby favoring nucleation of a buckled domain at the defect site and exhibiting rebuckling transition. Our prediction that rebuckling does not occur for small defects (*ε* < 0.1) may be used in classifying various positioning sequences.

The possibility of a sequence-induced defect makes the relationship between *ε* and *n* (Fig. 10) less exact. Placing a base-pair mismatch of size *n* inside a positioning sequence is expected to generate a larger defect than placing it at a random location. Enhancement of defect-facilitated buckling for a positioning sequence with unpaired bases may be relevant for its *in vivo* mismatch repair. Similarly, occurrence of a sequence-induced defect near one of the DNA ends may favor plectoneme positioning at that end over the other [5, 57].

An alternate model, where the defect is associated with an increased bending energy cost may also be useful. *In vivo*, such a defect will disfavor in-situ nucleosome assembly, thus regulating genome access for DNA-binding proteins. Studying supercoiled DNA with multiple defects is an interesting future prospect. Such studies may show pinning of a plectoneme domain at a defect site after an unpinning from a larger defect. Furthermore, studying the role of defects on the mechanics of chromatin fibers may also be an interesting future possibility.

To conclude, we have presented a theoretical model for DNA buckling and the analyzed the consequences of introducing a defect. We provided an explicit theoretical treatment of thermal fluctuations in plectonemic DNA that may be relevant in modeling fluctuations of geometrically constrained polymers. We also classified defects depending on their size which leads to various possible states corresponding to different defect sizes that can be probed experimentally.

## Acknowledgments

Work at NU was supported by the National Institutes of Health (NIH) [Grants R01-GM105847, U54-CA193419 (CR-PS-OC), and a subcontract to grant U54-20

DK107980], and by the National Science Foundation (Grants MCB-1022117 and DMR-1206868). S.B. acknowledges support from a Molecular Biophysics Training Program at NU. Work at NIH was supported in part by the Intramural Research Program of the National Heart, Lung, and Blood Institute, National Institutes of Health.

## Appendix A Plectoneme Hamiltonian

A plectoneme structure made up of total DNA length *L_p_*, can be considered as two helices of length *L_p_/*2 wrapped around each other. The total Hamiltonian of the plectoneme structure is given by,

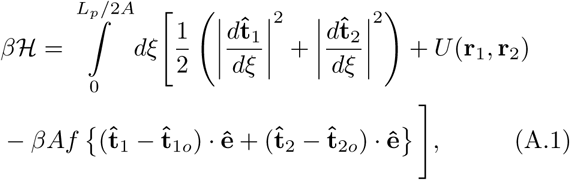

where an external force *f* is applied at the two ends of the plectoneme in a direction perpendicular to the axis of the plectonemic helix. This force can be interpreted as the stretching tension required at the end of a plectoneme that keeps the helices from unwinding themselves. Since, the force is in a direction perpendicular to the axis of the plectoneme, there is no corresponding force-extension energy contribution, however, the tension plays an important role in controlling the transverse fluctuations of the DNA inside the plectoneme.

In Eq. (A.1), the first parenthesized term containing the square of the local curvature of the two intertwining helices inside the plectoneme corresponds to the total elastic bending energy. The second term contains the total electrostatic energy contribution from close proximity of the DNAs inside the plectonemic structure [4, 21, 34, 58]. And, the third term contains the coupling of the transverse fluctuations of the DNAs to the external force, where ê is the direction of the plectoneme axis.

The above Hamiltonian is similar to the one analyzed for two intertwined DNAs or braids in Ref. [34], except for that braids have a force-extension energy [34], while plectonemes do not. Note in Eq. (A.1), the external force is only coupled to the transverse tangents.

### A.1. Oscillating Reference Frame

We consider two sets of orthonormal triads: 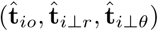, where *i* ∈ {1,2} corresponds to the two intertwining helices in the plectoneme. 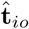 points along an average helical shape defined by two helix parameters: radius (*r*) and pitch *(2*π*p);* 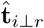 points towards the axis of the plectoneme; whereas 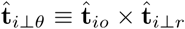. See Ref. [34] for a similar calculation done in the context of intertwined DNAs or DNA braids.

**FIG. 13.**
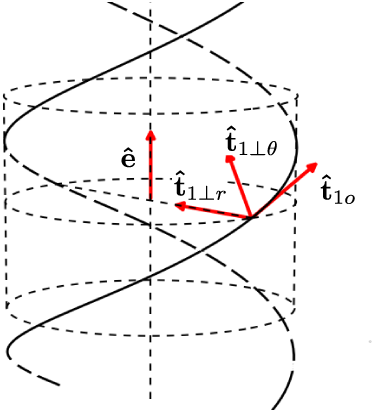
Schematic of the orthonormal triad 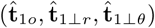 where **ê** shows the axis of the plectoneme superhelix.

We expand the tangent vectors to harmonic order, about a mean helical shape 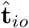 [Fig. (13)]:

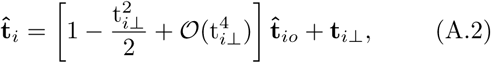

where the mean helical shape depends on two helix parameters radius (*r*) and pitch *(2*π*p):*

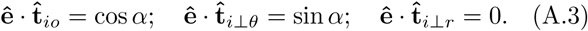

Here, *α =* arctan(r/p), is the braiding angle.

### A.2. Electrostatic Interactions

The electrostatic energy contribution due to DNA-DNA repulsion in a helical structure <scU> computed from a Debye-Hu¨ckel-type interaction is as follows.

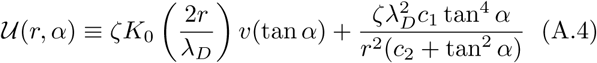

The first term in Eq. (A.4) is the contribution from electrostatic interaction with the neighboring strand in the superhelix. Here, *K*_0_(*x*), the modified Bessel function of the first kind, denotes the solution for two parallel strands [4, 58], and *v*(*x*) *=* 1 + (0.828)*x*^2^ + (0.864)*x*^4^, is an enhancement factor that accounts for the effect of helical curvature [21]. The second term accounts for the self interaction of the helically-bent polymer in the plectoneme, where *c*_1_ *=* 0.042 and *c*_2_ *=* 0.312 [34]. λ*_D_* is the Debye length of the ionic solution, and we define ζ = *2Aℓ_b_ν^2^*, a parameter that depends on the effective linear charge density of DNA *ν* [21, 44, 59, 60], and ℓ*b ≈* 0.7 nm is the Bjerrum length of water at *290 K.* We have used *ν =* 1.97, 6.24, 8.85, 10.23, and 26.6 nm^-1^ respectively corresponding to 0.01, 0.1, 0.15, 0.2, and 0.5 M monovalent salt concentrations [21, 23, 34, 44].

We define the electrostatic part of the Hamiltonian as: 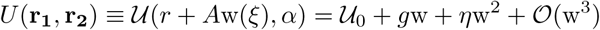 where 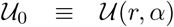; 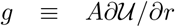, and 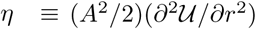 is the effective modulus of the electrostatic potential. The first term gives the mean electrostatic energy per unit length *A* of plectoneme with fixed radius and pitch, while the subsequent terms are corrections for small uniform deviation in the braid radius. Here, **w**(*ξ*) is the normalized radial deformation, defined as:

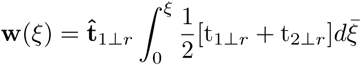

where tij__r_ are given by Eq. (A.2) and we assume the boundary condition w(0) = 0. Note, the above definition of normalized radial deformations w(ξ) assumes a parallel configuration of the two strands.

### A.3. Thermal Fluctuations

#### a. Perturbative expansion of the Hamiltonian

Following the above equations, the plectoneme Hamiltonian [Eq. (A.1)] can be expanded as a contribution from the mean-field helix structure (<scH>_0_) and thermal fluctuations (Δ<scH>) [34]:

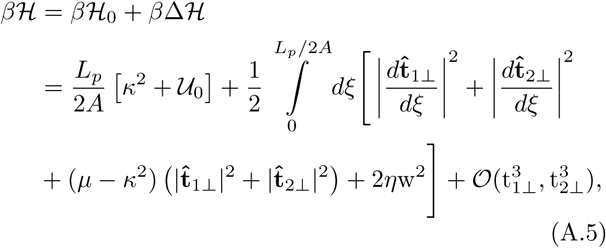

where *µ* ≡ *βAf cos α*, is the effective tension in the plectoneme; κ ≡ *Ar/(r^2^* +*p^2^)*, is the total mean-field curvature per unit persistence length of the strands. The mean-field term, (<scH>0) is the sum total of bending and electrostatic energy in the plectoneme [Eqs. (B.1) and (B.7)]. The sub-leading order term *(∆*<scH>) is the contribution from thermal fluctuations. We set the reference of the fluctuation free energy by setting the amplitude of the zero-momentum mode of transverse fluctuations to zero [34].

*b. Free energy of fluctuations* We construct a partition function for the plectoneme via a path integral over all the transverse tangent conformations:

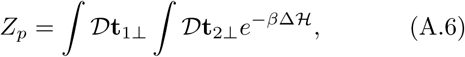

and get the free energy contribution of thermal fluctuations from the partition function [34],

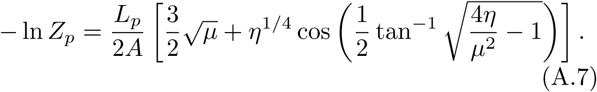

The first term gives the fluctuation free energy that depends only on the external tension, whereas, the second term depends on both the external tension and the salt concentration of the solution.

Note that there are four independent degrees of transverse fluctuations in a plectoneme structure: 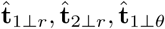, and 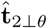 [Fig. (13a)]. Three of them 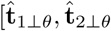, and 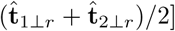 are solely controlled by the tension µ; while, the fluctuations in the direction 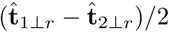 are controlled by both the external tension µ and the salt concentration via the parameter η. We have used the above expression for the fluctuation free energy inside the plectoneme structure in Eq. (5).

*c. Radial fluctuations.* Fluctuations in the radius of the plectoneme are generated by displacement of one plectonemic strand relative to the other. As is the case for Gaussian fluctuations, the two-point correlation function of radial deformations decays exponentially: 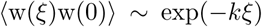, where 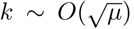 [34]. The zero-distance behavior of the two-point correlation gives the radial fluctuations in the plectonemic superhelix (See Appendix B in Ref. [34]):

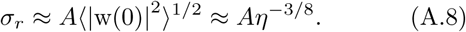

This suggests that a stronger electrostatic repulsion reduces fluctuations in the plectoneme radius.

## Appendix B: Finite-Sized Supercoiled DNA

### B.1. Defect-free DNA

#### a. Buckled state: plectonemes and end loops

The total free energy of the buckled state composed of *m* domains of plectoneme:

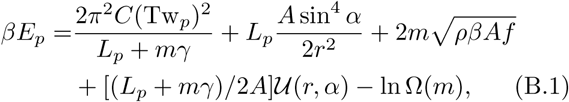

where the first term corresponds to DNA-twist energy contribution in the buckled state. The second and third terms respectively correspond to the net elastic energy of plectoneme superhelices and m end loops. The fourth term contains the total mean-field electrostatic contribution from the buckled state of the DNA. Finally, the last term in Eq. (B.1) corresponds to configuration entropy of *m* plectoneme domains (m ≥ 1) [23, 34], where

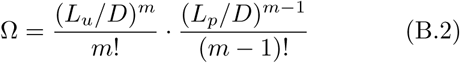

is the total number of energetically degenerate but distinguishable configurations arising from: (1) one-dimensional diffusion of a domain along the DNA contour length [the first combinatorial term in Eq. (B.2)]; and (2) the exchange of DNA length among the plectoneme domains [the second combinatorial term in Eq. (B.2)]. 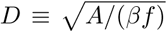, is the force-induced correlation length, which we have used as the distinguishable length for plectoneme sliding.

#### b. Force-extended state

The total energy of the force-coupled state is given by

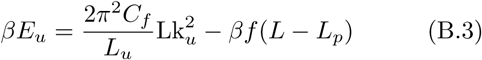

The first term corresponds to the total twist energy, where 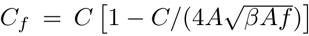 is the renormalized twist persistence length [20]. The second term is the extension energy of the DNA under external force f.

#### c. Thermal fluctuations

The total fluctuation contribution is obtained from summing the contributions from the force-extended and plectoneme states:

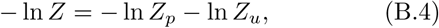

where *Z_p_* corresponds to the plectoneme [Eq. (A.7)]; and *Z_u_* corresponds to the force-extended state, which is computed by taking the *η* → 0 limit of Eq. (A.6). *–*ln *Z_u_* is the second term in Eq. (5).

#### d. Numerical scheme

Various quantities of interest can be numerically computed from the partition function *Z* in Eq. (6) as follows:

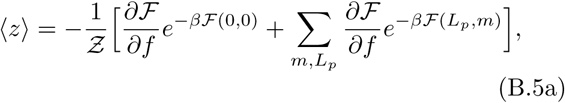

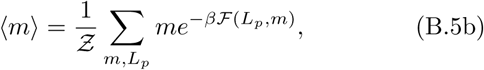

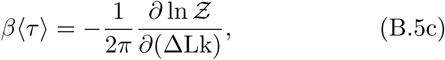

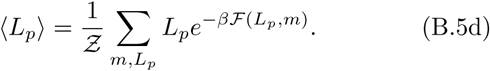

#### e. Probability distributions

For a given coexistence state *(L_p_,m)*, the probability distribution of *X* ∈ {*z,τ*} is given by

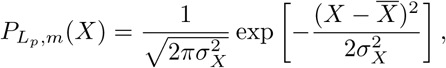

where the mean 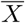 and the standard deviation σ_X_ are obtained as follows:

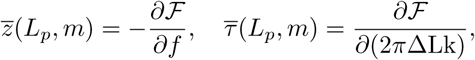

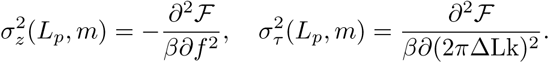

Now, the total probability distribution of *X* for a given linking number and force is obtained by summing the contributions from all the states considered in the partition sum:

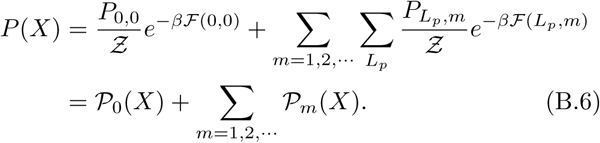

In the above equation, the probability distribution is written as a sum of contributions from the force-extended state, and the buckled domain containing *m* end loops.

### B.2. DNA with a Defect

#### a. Free energy of the buckled state

The free energy of the plectoneme state, now including m mobile plectoneme domains and *m*^†^ pinned plectoneme domains, is given by

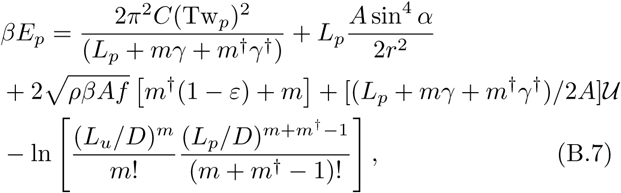

where the first and the second terms respectively correspond to the total twist and bending energy in the plectoneme state. The third term gives the total elastic energy associated with *m* mobile and *m*^†^ pinned plectoneme end loops. The fourth term corresponds to net electrostatic energy of the buckled state. And the fifth term is associated with the total configuration entropy of plectoneme domains.

*Theta function.*

We have used the usual definition of Theta function in the partition sum [Eq. (12)].

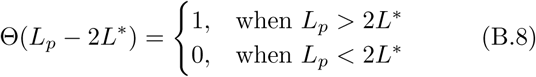

*b. Probability distributions.* Similar to Eq. (B.6), we write the probability distribution of *X* at a fixed linking number and fixed force as:

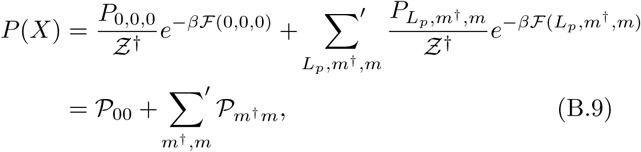

where the sum is now a restricted one as shown in the partition sum [Eq. (12)]. The contribution from the buckled state with m and m’ mobile and pinned plectonemes respectively is 𝒫*_m_*_†_*_m_.*

